# Multiple Determinants and Consequences of Cohesion Fatigue in Mammalian Cells

**DOI:** 10.1101/240630

**Authors:** Hem Sapkota, Emilia Wasiak, John R. Daum, Gary J. Gorbsky

## Abstract

Cells delayed in metaphase with intact mitotic spindles undergo cohesion fatigue, where sister chromatids separate asynchronously, while cells remain in mitosis. Cohesion fatigue requires release of sister chromatid cohesion. However, the pathways that breach sister chromatid cohesion during cohesion fatigue remain unknown. Using moderate-salt buffers to remove loosely bound chromatin Cohesin, we show that “cohesive” Cohesin is not released during chromatid separation during cohesion fatigue. Using a regulated protein heterodimerization system to lock different Cohesin ring interfaces at specific times in mitosis, we show that the Wapl-mediated pathway of Cohesin release is not required for cohesion fatigue. By manipulating microtubule stability and Cohesin complex integrity in cell lines with varying sensitivity to cohesion fatigue, we show that rates of cohesion fatigue reflect a dynamic balance between spindle pulling forces and resistance to separation by interchromatid cohesion. Finally, while massive separation of chromatids in cohesion fatigue likely produces inviable cell progeny, we find that short metaphase delays, leading to partial chromatid separation, predispose cells to chromosome missegregation. Thus, complete separation of one or a few chromosomes and/or partial separation of sister chromatids may be an unrecognized but common source of chromosome instability that perpetuates the evolution of malignant cells in cancer.

## Introduction

Cells delayed or arrested at metaphase with intact mitotic spindles undergo cohesion fatigue, where sister chromatids separate asynchronously, while the cells remain in M phase (Daum et al., 2011; Stevens et al., 2011). Separated chromatids generated before anaphase likely missegregate or form merotelic attachments that can result in aneuploidy and chromosome breakage. While all cells can undergo cohesion fatigue when arrested at metaphase, the rate of chromatid separation varies significantly within a population of cells and among different cell types, even those closely related. Microtubule pulling forces are essential. Treatment of cells with Nocodazole, a microtubule depolymerizer, completely eliminates cohesion fatigue in mitotic cells arrested by treatment with the proteasome inhibitor, MG132, or by depletion of the SKA3 protein (Daum et al., 2011).

The Cohesin complex normally holds sister chromatids together from DNA replication until anaphase (Michaelis et al., 1997). The major structural elements of the Cohesin ring consists of two Structural Maintenance of Chromosome proteins (SMC3 and SMC1) and cohesin complex component RAD21 that closes the ring. These proteins intersect at three sites, referred to as “gates.” Cohesin gates may open during different stages of dynamic Cohesin-chromatin interactions during the cell cycle. For example, Cohesin appears to load onto chromosomes via the opening of the SMC3 and SMC1 hinge interface (Buheitel and Stemmann, 2013) and partially through the SMC3 and RAD21 interface (Murayama and Uhlmann, 2015). To release sister chromatids from each other in mitosis in vertebrates, Cohesin complexes are removed from chromosomes through two mechanisms. In early mitosis until metaphase, the ‘prophase pathway’ uses Plk1 and Aurora B kinases and the Cohesin removal protein, Wapl, to release a large portion of Cohesin from chromosome arms via opening of SMC3-RAD21 interface of Cohesin. Then at the metaphase-anaphase transition, the protease, Separase, cleaves the RAD21 component of the remaining chromosome-bound Cohesin to induce the final separation of sister chromatids (Waizenegger et al., 2000).

In addition to its three core structural ring components, the Cohesin complex contains several regulatory, auxiliary components. One of these has two isoforms called Stromal Antigen 1 and 2 (SA1 or SA2) (Solomon et al., 2011; Sumara et al., 2000). Cohesin complexes contain either SA1 or SA2 (Zhang et al., 2008). Cells depleted of either SA1 or SA2 continue to proliferate, but deletion of both is lethal (van der Lelij et al., 2017). Cohesin complexes containing SA1 appear important for arm and telomere cohesion, while Cohesin complexes containing SA2 have more critical roles for centromeric cohesion (Canudas and Smith, 2009). SA2 at centromeres recruits proteins that promote cohesion, including Sororin, Shugoshin (SGO1), and Protein Phosphatase 2A (PP2A), that shield centromeric Cohesin from phosphorylation and removal due to the Wapl-mediated prophase pathway (Hauf et al., 2005; McGuinness et al., 2005; Nishiyama et al., 2013).

The separation of chromatids in cohesion fatigue requires release of sister chromatid cohesion. However, we do not know if and how the Cohesin complex is breached during cohesion fatigue. Although we and others have shown that depletion of Wapl, a negative regulator of cohesin, prior to mitotic entry, delays cohesion fatigue, it is unclear whether continued Wapl activity is essential for cohesion fatigue after the chromosomes align at the metaphase plate. Previously, we reported that Cohesin protein levels in chromosome fractions remained constant before and after cohesion fatigue (Daum et al., 2011). However, a subsequent study indicated that most, but not all Cohesin in isolated chromosomes was released by a treatment with a moderate concentration of salt (Bermudez et al., 2012). This result suggested that the only the minor, the salt-resistant population comprises the “cohesive” Cohesin that functionally holds sister chromatids together. Currently, we do not comprehensively understand the factors that determine the sensitivity of cells to cohesion fatigue, the mechanism by which cohesion is lost during fatigue, and the consequences of partial and full chromatid separation to downstream chromosome instability.

## Results

### Cohesin remains bound to chromatids after fatigue

Current models indicate that Cohesin is released from chromosomes during chromatid separation at anaphase (Supplementary figure 2A and (Kueng et al., 2006; Tomonaga et al., 2000; Uhlmann et al., 2000) Cohesion fatigue also generates separated chromatids. Thus we anticipated that Cohesin should also be released from chromosomes during the process. Nevertheless, in our previous work comparing isolated chromosomes and chromatids prepared from cells before and after fatigue, surprisingly, levels of the core Cohesin subunits associated with chromatin remained unchanged (Daum et al., 2011). However, a potential explanation for this result came from a subsequent study, which revealed that in isolated mitotic chromosomes most Cohesin can be released by treatment with moderate salt (Bermudez et al., 2012). The implication of that work was that only the minor, salt resistant Cohesin was functional in sister chromatid cohesion, and perhaps this small pool was indeed released during fatigue but was too small for detection in our previous study.

We first confirmed that only a fraction of Cohesin remains bound to chromosomes after treatment with moderate salt buffer (Supplementary figure 1B). We then examined whether any changes occurred in the salt-resistant population before and after cohesion fatigue. We treated mitotic cells with MG132 in the absence or presence of Nocodazole for 8 h and then isolated chromosome fractions in moderate salt buffer. As expected, more than 90% of cells treated with MG132 without Nocodazole showed over half of their chromatids separated compared with only 5% of cells treated with MG132 in the presence of Nocodazole. If the salt-resistant Cohesin was released during fatigue, at least a 45% reduction (dotted line figure 1B) in core Cohesin should occur in fatigued samples (MG132 alone) compared to non-fatigued samples (MG132 + Nocodazole). However, immunoblotting for the core Cohesin component, SMC3 revealed no differences Cohesin levels between fatigued and non-fatigued samples (Figures 1A and 1B). Thus, Cohesin release did not occur during cohesion fatigue tracking either the total chromosome-bound population or the salt-resistant population.

**Figure 1:**
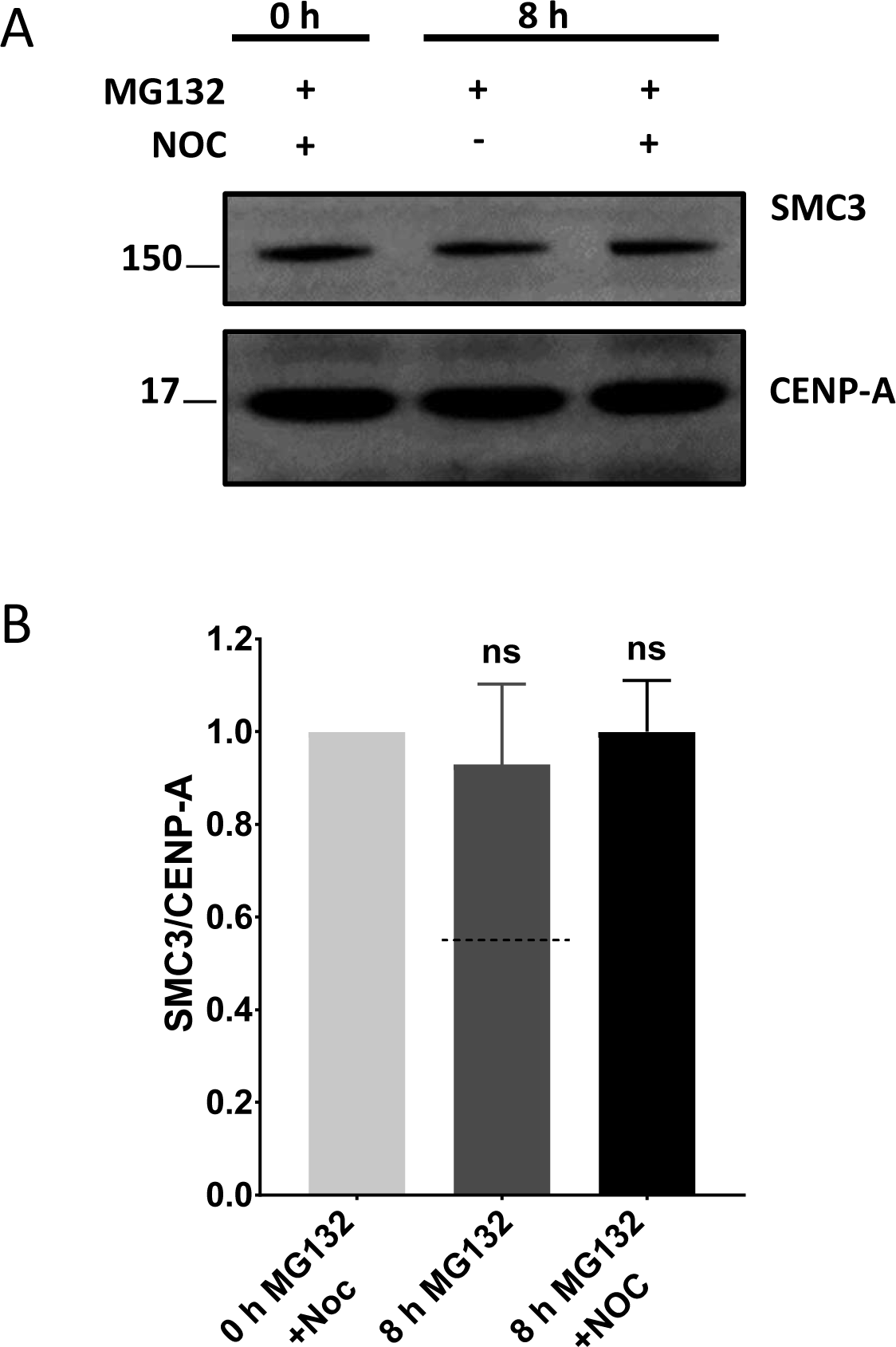
The core Cohesin protein SMC3 remains bound to chromatids during cohesion fatigue. **(A)** Immunoblotting of chromosome fractions from mitotic HeLa cells treated with MG132 ±Nocodazole for 8 h. Lane 1 shows baseline salt-resistant SMC3 level in mitotic chromosomes before cohesion fatigue. Lane 2 reflects SMC3 in chromosome fractions of fatigued chromatids. Lane 3 shows SMC3 in negative control for cohesion fatigue (MG132 +Nocodazole). **(B)** Quantified immunoblots from four independent experiments where band intensity of SMC3 was measured and normalized to CENP-A band intensity. Dotted line on graph represents the expected level of SMC3 if cohesin was lost from fatigued chromosomes based on the percentage of separated chromosomes seen in chromosome spreads. Kruskal-Wallis test with Dunn’s multiple comparison was used for statistical analysis. Error bars represent standard deviation.

### Centromeric levels of Shugoshin1 are not critical regulators of sensitivity to cohesion fatigue

The Shugoshin1 (SGO1) protein protects centromeric Cohesin from Wapl-mediated release by recruiting protein phosphatase 2A (PP2A) to the centromere region (Gandhi et al., 2006; Liu et al., 2013b; Shintomi and Hirano, 2009; Xu et al., 2009). Changes in SGO1 have been implicated in explaining why different cell lines show differential sensitivity to cohesion fatigue during metaphase arrest (Liu et al., 2013a; Tanno et al., 2015). For our studies, we used two isolates of HeLa cells that exhibit strong differences in the rate of cohesion fatigue (Supplementary figures 4A and 4B). One HeLa cell line, stably expressing histone H2B-GFP, undergoes cohesion fatigue with an average time of approximately 340±127 min at metaphase, while another HeLa cell line, stably expressing histone H2B-mRFP, undergoes cohesion fatigue after an average of 130±55 min. We named these cell lines HeLa-Slow and HeLa-Fast, respectively. We induced metaphase arrest by treating cells with MG132 or with ProTAME, a cell permeant inhibitor of the Anaphase Promoting Complex/Cyclosome (APC/C) (Lara-Gonzalez and Taylor, 2012; Sackton et al., 2014; Zeng et al., 2010).

We measured total SGO1 levels in both HeLa-Fast and HeLa-Slow cells and compared SGO1 levels by immunofluorescence in normal prophase, prometaphase or metaphase cells and in cells arrested in metaphase for 6 h (HeLa-Slow) or 3 h (HeLa-Fast). From metaphase-arrested cells, we selected fatigued cells and examined their SGO1 levels. In HeLa-Slow cells, SGO1 levels diminished from prometaphase to metaphase, but no further reduction in SGO1 levels occurred with metaphase arrest for 6 h. Cells with separated chromatids showed no reduction in SGO1 levels compared to normal metaphase-arrested cells (Supplementary figure 1D). HeLa-Fast cells showed a similar trend during mitotic progression with SGO1 showing reduced levels at metaphase. In these cells, SGO1 levels were further decreased after 3 h of metaphase delay. However, in cells that underwent cohesion fatigue during the 3 h metaphase delay, SGO1 levels were equal to levels of normal metaphase cells (Supplementary figures 1E). Thus, both cell lines showed a reduction in centromere-associated SGO1 levels as the cells aligned their chromosomes, but SGO1 did not appear to be altered during fatigue. Finally, comparison of total chromosome associated SGO1 levels showed higher levels in HeLa-Fast cells than in HeLa-Slow cells, the opposite that might be expected if SGO1 levels were a major determinant of resistance to cohesion fatigue (Supplementary figures 1C).

### Inhibiting Wapl-mediated Cohesin release during early mitosis delays subsequent cohesion fatigue

We and others have previously shown that depletion of Wapl, which mediates Cohesin removal during early mitosis, delays cohesion fatigue (Daum et al., 2011; Stevens et al., 2011). To extend these studies in a system where enhanced Cohesin binding to mitotic chromosomes could be directly monitored, we used Hela cells stably expressing SMC1-GFP (Hou et al., 2007) and examined the effects of Wapl depletion. We depleted Wapl via RNAi, treated the SMC1-GFP cells with ProTAME, and examined cells with clear SMC1-GFP signals on metaphase chromosomes, indicative of those with efficient Wapl depletion (Supplementary figures 2A). Normally, Cohesin released into the cytoplasm by the Wapl-mediated prophase pathway obscures the residual chromosome-bound population, but Wapl depletion results in strong retention of chromosome Cohesin (Gandhi et al., 2006; Haarhuis et al., 2013; Haarhuis et al., 2017; Tedeschi et al., 2013). When Wapl-depleted cells were arrested at metaphase, there was a significant increase in time these cells take to undergo cohesion fatigue (Figure 2A), confirming that Wapl depletion causes increased chromosome association of Cohesin that in turn delayed cohesion fatigue without affecting the total number of cells undergoing fatigue. As another approach we manipulated a competitor of Wapl activity, Sororin, which is normally released from chromatin by mitotic phosphorylation. A Sororin mutant (9A-Sororin) resists mitotic phosphorylation and inhibits Wapl-mediated Cohesin release (Liu et al., 2013b). As expected, cells expressing the 9A mutant form of Sororin showed delayed cohesion fatigue compared to cells expressing wild type Sororin (Figure 2B).

**Figure 2:**
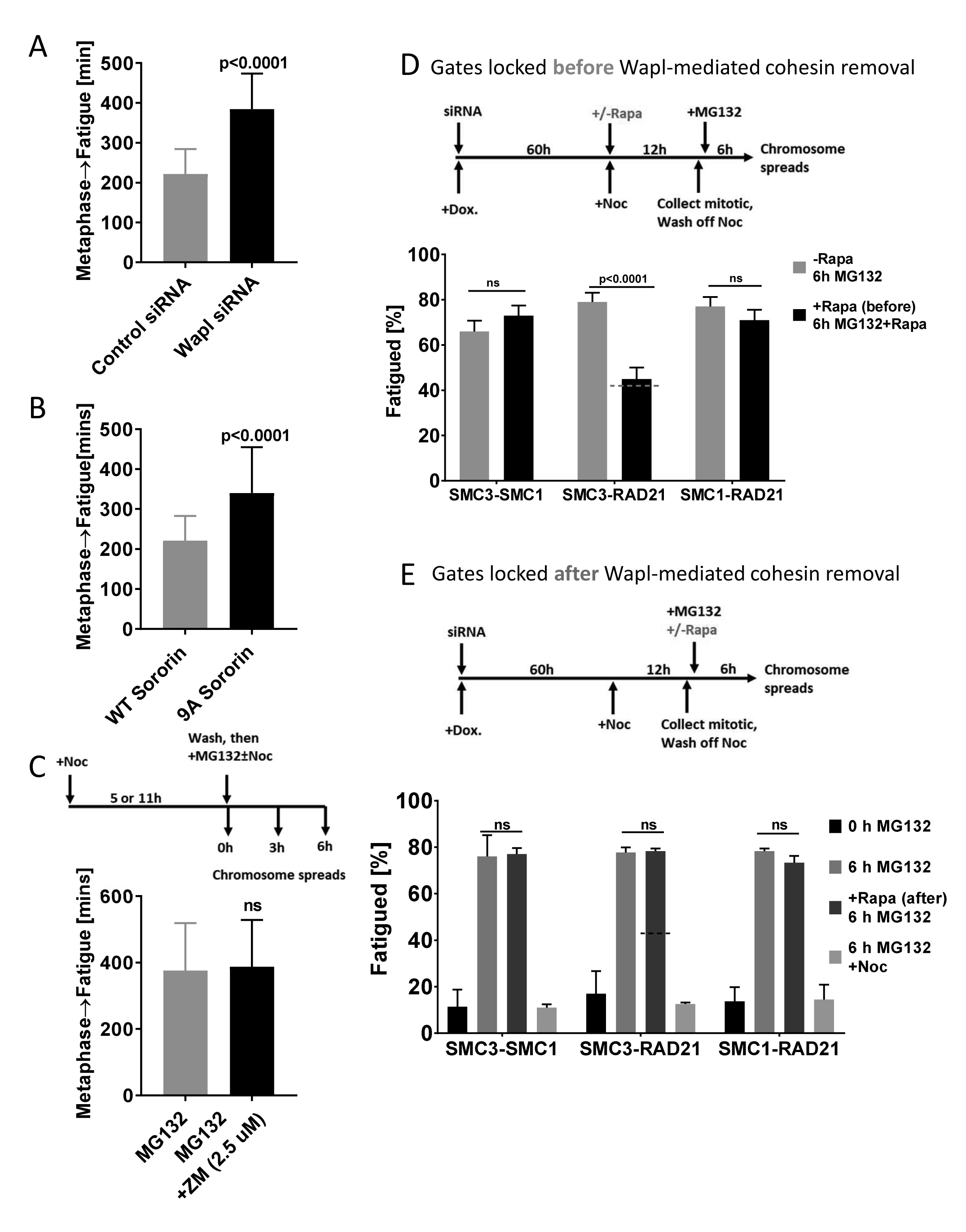
Wapl-mediated release of Cohesin after metaphase is not required for cohesion fatigue. **(A)** Wapl depletion during interphase slows cohesion fatigue in subsequent mitosis. Elapsed times from metaphase to chromatid separation/cohesion fatigue were determined via live cell imaging in HeLa cells stably expressing SMC1-GFP. In Wapl-depleted cells, only cells with clear GFP signal on chromosomes at the metaphase plate (indicating successful Wapl depletion) were scored in two independent experiments with totals of >100 cells. **(B)** Expression of phosphorylation resistant Sororin slows cohesion fatigue. Elapsed times from metaphase to cohesion fatigue were determined in HeLa cells expressing either WT sororin or non-phosphorylatable 9A-sororin. At least 60 cells were scored for each cell type. The Mann-Whitney test was used for statistical analysis. **(C)** Inhibition of Aurora B kinase after metaphase alignment does not inhibit cohesion fatigue. Experimental scheme and graph depicting elapsed times from metaphase to cohesion fatigue were determined after 2.5 uM ZM 447439 treatment in cells released from Nocodazole to MG132 for 1 h. Three independent experiments with totals of >200 cells were quantified. The Mann-Whitney test was used for statistical analysis. **(D)** Locking the SMC3-Rad21 gate but not other gates before mitotic entry inhibits cohesion fatigue. Experimental scheme and results from chromosome spreads in Hek293 expressing Cohesin fusions to Rapamycin-binding proteins treated with Rapamycin to lock specific gates **before cells entered mitosis** then treated with MG132 for 6 h to arrest cells at metaphase and allow cohesion fatigue. Totals of >100 spreads per condition per cell line were quantified. Graph shows mean±S.E.M. Dotted line represents the expected inhibition of fatigue with efficient SMC3-RAD21 gate locking based on % of spreads with unresolved chromatid arms (45%) from supplementary figure 2D. **(E)** Locking any of the Cohesin gates after completion of Wapl-mediated Cohesin release in early mitosis does not inhibit cohesion fatigue. HEK293 cells expressing Cohesin fusions to Rapamycin-binding proteins were treated with or without Rapamycin **after allowing completion of early mitosis, Wapl-mediated Cohesin removal** in three independent experiments with totals of > 450 spreads per cells line for each treatment. Graph shows Mean±S.D. Two-way ANOVA with Tukey’s multiple comparison test was used for statistical analysis.

### Inhibiting the Wapl pathway or locking cohesin gates after chromosome alignment at metaphase does not inhibit cohesion fatigue

The studies above show that inhibiting the Wapl-mediated Cohesin release during early mitosis slowed cohesion fatigue. Inhibition of Wapl function before mitotic entry increased the levels of salt-resistant Cohesin retained on chromosomes (Supplementary figures 2A), and this might fully account for delays in cohesion fatigue. However, it remained possible that Wapl continues to function in opening Cohesin rings during metaphase arrest, contributing to cohesion fatigue after chromosome alignment. We used two distinct approaches to test this possibility. Two mitotic kinases, Plk1 and Aurora B are critical for the function of the Wapl. In our previous work, we showed that chemical inhibition of Plk1 did not block cohesion fatigue, but in that study, the rates of fatigue were not quantified (Daum et al., 2011). Here we used ZM447439, an inhibitor of Aurora B kinase to inhibit Wapl-mediated Cohesin release in HeLa-Slow cells after chromosome alignment at metaphase. Treatment of these cells in early mitosis with 0.5 µM ZM447439 caused significant defects in chromosome alignment, confirming inhibition of Aurora B kinase (Supplementary figures 2B). We added the inhibitor at 2.5 µM, a fivefold higher concentration, one hour after release from Nocodazole to MG132 after most cells had aligned their chromosomes, to avoid disrupting chromosome alignment. Then we tracked cells with tight metaphase plates at the time of ZM447439 addition. The addition of 2.5 µM ZM447439 did not induce loss of chromosome alignment in cells at metaphase, and did not delay cohesion fatigue (Figure 2C). This finding indicated that the continued activity of Aurora B kinase in promoting Wapl activity in metaphase cells does not promote cohesion fatigue in these cells.

The Wapl-mediated prophase pathway releases Cohesin by opening the SMC3-RAD21 interface or gate. As a stringent test of the role of the Wapl in cohesion fatigue we used three HEK293 cells lines, each expressing a pair of Cohesin ring components tagged with FRB or FKBP proteins that allows locking of SMC3-RAD21 gate, the SMC1-RAD21 gate, and the SMC1-SMC3 gate by the addition of Rapamycin (Buheitel and Stemmann, 2013). We depleted endogenous Cohesin proteins and induced expression of the siRNA-resistant fusion proteins. We used chromosome spreads to confirm previously published work that locking the SMC3-RAD21 gate, but not the other gates in early mitosis, inhibited Wapl-mediated release of Cohesin and increased the proportion of chromosomes with unresolved chromosome arms (Supplementary figures 2C and D). We then studied the effect on cohesion fatigue (Figures 2D and E). As expected, when we locked the Cohesin gates by adding rapamycin before cells entered mitosis (Figure 2D top), chromosome spreads showed that cohesion fatigue was inhibited in cells expressing the SMC3-RAD21 pair of Rapamycin-binding proteins, mimicking Wapl inhibition (Figure 2D bottom). Locking the other two gates showed no effect on cohesion fatigue assayed with chromosome spreads. Identical results were found by live cell imaging (Supplementary figures 2E). The results obtained by locking gates before entry into mitosis reveal that gate locking was efficient, that locking the SMC3-RAD21 gate mimicked Wapl depletion in blocking the cohesin removal during prophase and prometaphase, and that Rapamycin-induced dimerization of SMC3-RAD21 in the presence of Wapl was robust and could resist chromatid separation by spindle-pulling forces.

Next, we locked the Cohesin gates of Cohesin on chromosomes in metaphase after the normal Wapl-mediated release of unprotected Cohesin during prophase and prometaphase. To accomplish this, we first incubated mitotic cells with nocodazole for 12 h to allow Wapl-mediated cohesin release to be completed (Figure 2E top). We then released cells from nocodazole to MG132 or MG132 plus nocodazole and added Rapamycin to lock each gate. Consistent with our results from Aurora B inhibition, chromosome spreads from cells treated for 6 h with MG132 showed that locking any of the Cohesin gates after metaphase alignment did not inhibit cohesion fatigue (Figure 2E bottom). These results indicate that inhibition of the Wapl before/early in mitosis delays cohesion fatigue through an increase in the amount of functional Cohesin retained on chromosomes. However, once cells are at metaphase, after full activity of the Wapl-mediated prophase pathway is complete, inhibition of the Wapl-mediated prophase pathway does not delay fatigue, indicating it is not required for chromatid separation. In addition, locking the other Cohesin gates does not affect fatigue. Thus, transient opening of a single Cohesin gate is unlikely to account for the separation of sister chromatids in cohesion fatigue for cells we have analyzed.

### Compromised Cohesin accelerates cohesion fatigue

Cohesin-chromatin interactions are highly regulated throughout cell cycle (Bermudez et al., 2012; Gandhi et al., 2006; Lara-Gonzalez and Taylor, 2012; Liu et al., 2013a; Whelan et al., 2012; Xu et al., 2014). In early mitosis, Cohesin is removed from chromosome arms by the Wapl-mediated prophase pathway (Gandhi et al., 2006; Kueng et al., 2006; Nishiyama et al., 2010; Nishiyama et al., 2013; Shintomi and Hirano, 2009). In cells arrested in mitosis for long periods, Cohesin removal separates chromosome arms, which generates the classic “X-shape” chromosomes seen in chromosome spreads (Supplementary figures 3A). We previously found that under normal conditions, cohesion fatigue initiates at kinetochores then propagates down the chromosome arms. Thus, Cohesin loss and arm separation should increase cell susceptibility to cohesion fatigue. To test this idea, we used chromosome spreads to compare rates of cohesion fatigue in LLC-PK cells after arresting cells in mitosis for 5 or 11 h with Nocodazole. After Nocodazole arrest, cells were washed then placed in fresh media containing MG132 and then processed for chromosome spreads immediately (0 h), 3 h or 6 h later (Figure 3A left). Cells arrested in Nocodazole for 11 h had significantly increased cohesion fatigue compared to cells arrested for just 5 h (Figure 3A right). In contrast, cells harvested at 0 h, 3 h or 6 h after being maintained in MG132 plus Nocodazole showed very few separated chromatids. We hypothesized that arrest in Nocodazole might decrease the level of salt-resistant Cohesin on chromosomes. Quantification of western blots showed a modest decrease in chromosome-bound Cohesin levels comparing chromosomes from cells arrested for 5 and 11 h (Supplementary figures 3C). These results indicate that longer mitotic arrest, without spindle pulling forces, primes cells to undergo faster cohesion fatigue.

**Figure 3:**
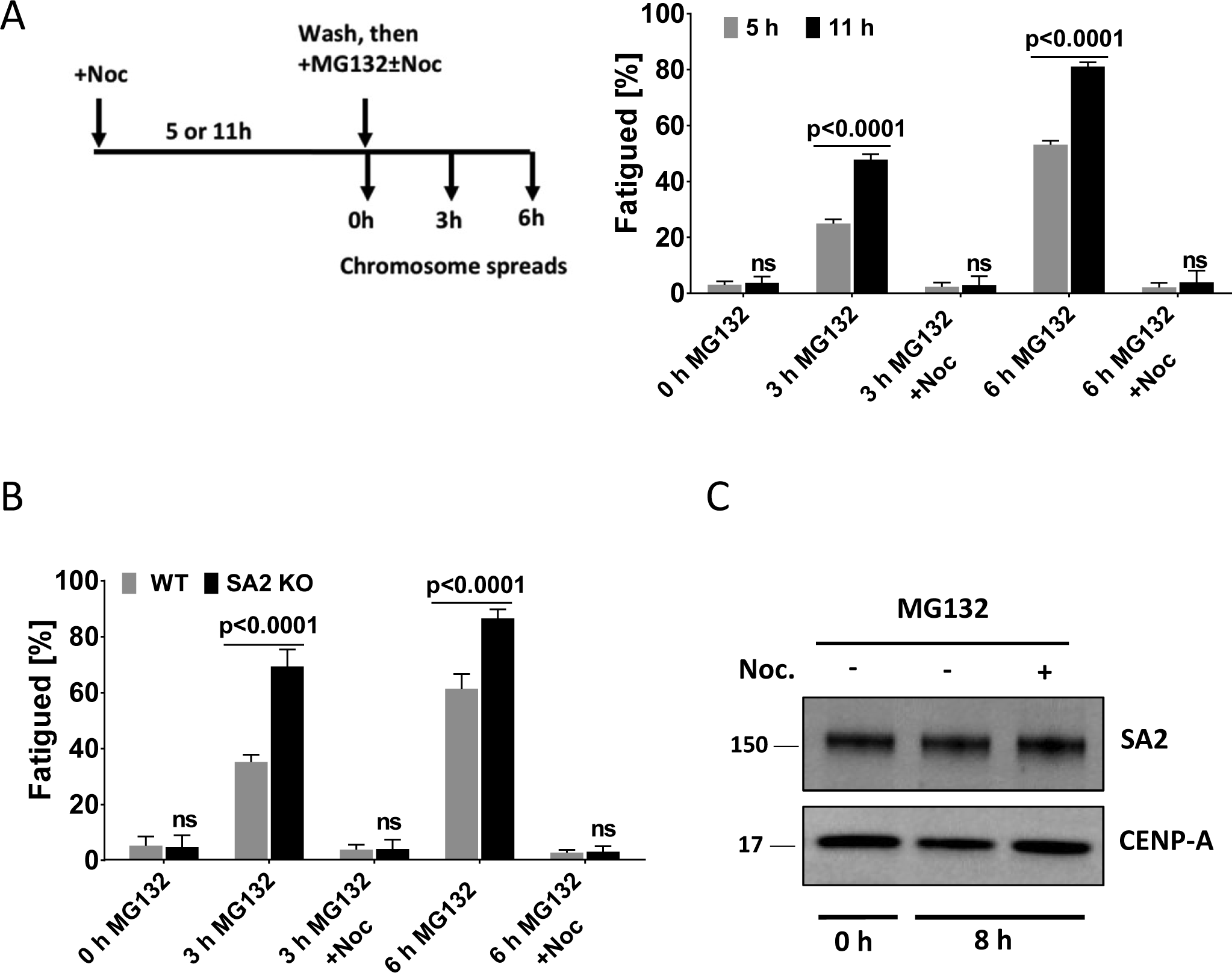
Altering Cohesin changes the rate of cohesion fatigue. **(A)** Longer mitotic arrest in Nocodazole leads to enhanced cohesion fatigue. Experimental scheme and chromosome spread analysis for LLC-PK cells arrested in mitosis for 5 or 11 h with 330 nM Nocodazole, then washed and treated with MG132 ±Nocodazole (330 nM) to arrest at metaphase for 3 or 6 h. Three independent experiments with a total of >600 spreads were scored for each treatment. Cells initially arrested for 11 h in Nocodazole undergo more rapid cohesion fatigue than those arrested for 5 h, consistent with the continued action of the Wapl-mediated prophase pathway in removing Cohesin from chromosomes in cells arrested in mitosis. Error bars show standard deviation. Two-way ANOVA with Sidak’s multiple test was used for statistical analysis. **(B)** SA2 knockout HCT116 cells undergo cohesion fatigue more rapidly than parental cells. Chromosome spreads were examined in parental and SA2 knockout HCT116 cells. Cells were treated with 330 nM Nocodazole overnight (16 h), then mitotic cells were collected, washed then treated with MG132 ±Nocodazole (330 nM) for 3 or 6 h. Three independent experiments with totals of >600 spreads were scored. Two-way ANOVA with Sidak’s multiple test was used for statistical analysis. **(C)** SA2 protein is not lost from chromatids during cohesion fatigue. Immunoblotting of SA2 protein in chromosome fractions prepared from mitotic HeLa cells treated with MG132 ±Nocodazole for 8 h. Lane 1 shows baseline SA2 levels in mitotic chromosomes, lane 2 shows SA2 in chromosome fractions from fatigued chromatids and lane 3 shows cohesion fatigue negative control (MG132 plus 330 nM Nocodazole for 8 h).

These results indicated that the increased time spent in mitosis leads to a higher propensity for cohesion fatigue. As a complimentary method to test this idea, we compared onset of cohesion fatigue in cells that reach full metaphase quickly with those where chromosome alignment is delayed. To increase the proportion of cells with alignment delays, we treated cells with 1.5 uM S-Trityl-L-cysteine (STLC), an inhibitor of mitotic motor kinesin Eg5 (Skoufias et al., 2006). When we measured the time from full metaphase alignment to cohesion fatigue, cells with the slowest alignment, and thus with longer times spent in prometaphase, showed faster cohesion fatigue (Supplementary figures 3B).

The Cohesin subunit SA2 is thought to promote cohesion specifically at centromeres (Canudas and Smith, 2009). Unlike Sgo1 depletion where sister chromatids separate without spindle pulling forces, depletion of SA2 caused increased interkinetochore distances only in the presence of intact spindles (Kleyman et al., 2014) suggesting defective cohesion maintenance rather than compromised cohesion establishment. If so, then depletion of SA2 should accelerate cohesion fatigue. We investigated the consequences of SA2 loss using HCT116 cells in which the STAG2 gene, which codes for SA2, had been deleted by homologous recombination (Solomon et al., 2011). Chromosomes from SA2 knockout cells showed reduced amounts of the Cohesin ring components, SMC3 and RAD21, compared to parental HCT116 cells (Supplementary figures 3E). Metaphase arrest for 3 or 6 h caused increased separation of chromatids in chromosome spreads of SA2 knockout cells compared to parental cells (Figure 3B). Inclusion of nocodazole to disrupt spindle microtubules abrogated the differences in chromatid separation in SA2 knockout and parental cells. Thus, loss of the Cohesin regulatory component, SA2, increases the susceptibility of cells to cohesion fatigue in the presence of spindle pulling forces. Because SA2 helps to resist cohesion fatigue, we hypothesized that its release might accompany fatigue. We analyzed chromosome fractions from HeLa cells by western blot before and after fatigue but found no reduction in the amount of chromosome-bound SA2 after chromatid separation (Figure 3C).

### Modulating microtubule stability alters rates of cohesion fatigue

Previously we showed that complete disruption of spindle microtubules blocked cohesion fatigue, which indicated that spindle pulling forces were essential (Daum et al, 2011). However, it was unclear the degree to which the rate of cohesion fatigue might be sensitive to microtubule dynamic turnover. To alter microtubule dynamics while maintaining intact spindles, we used 5 nM Nocodazole and 1.5 nM Taxol, concentrations that slow but do not block progression of control cells through mitosis (Supplementary figures 4C). We measured the elapsed time from metaphase to chromosome scattering (fatigue) in cells arrested at metaphase. Treatment with 5 nM Nocodazole marginally delayed chromosome alignment, but significantly slowed cohesion fatigue in both HeLa-Slow and HeLa-Fast cells (Figure 4A). Correspondingly, partial stabilization of spindle microtubules with 1.5 nM Taxol led to faster cohesion fatigue in both cell types (Figure 4B). When viewed as percentages, HeLa-Slow and HeLa-Fast cells showed comparable delay in cohesion fatigue when treated with Nocodazole and comparable acceleration when treated with Taxol.

**Figure 4:**
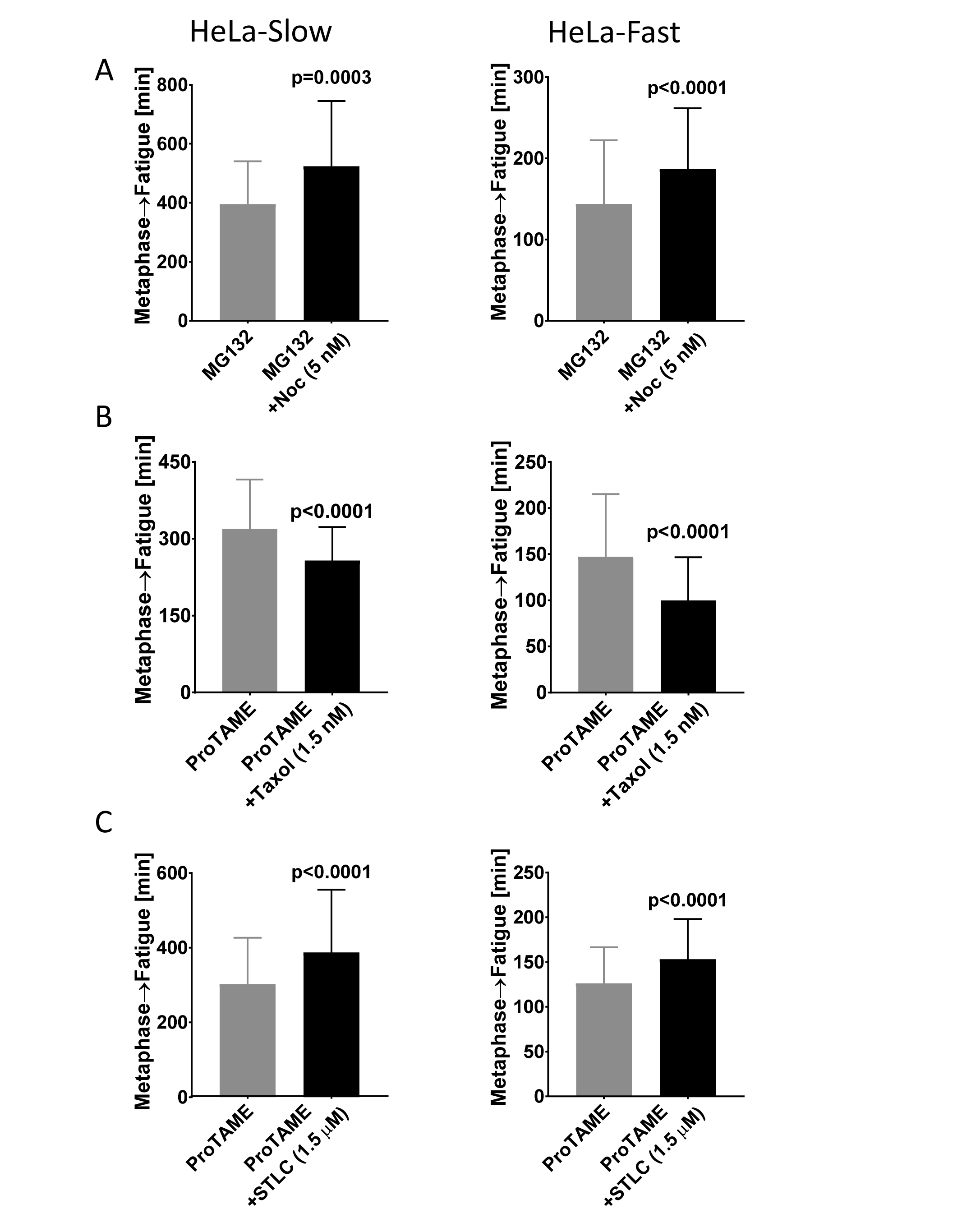
Microtubule dynamics and spindle tension impact cohesion fatigue. **(A)** Treatment of cells with low concentration of Nocodazole slows cohesion fatigue. The elapsed time from metaphase to chromatid scattering/cohesion fatigue was determined in HeLa-Slow cells (left) and HeLa-Fast cells (right) arrested at metaphase with MG132 +/- 5nM Nocodazole **(B)** Treatment of cells with low concentration of Taxol accelerates cohesion fatigue. The elapsed time from metaphase to chromatid scattering/cohesion fatigue was determined in HeLa-Slow cells (left) and HeLa-Fast cells (right) arrested at metaphase with ProTAME +/- 1.5nM Taxol. **(C)** Decreasing spindle tension with low concentration of STLC slows cohesion fatigue. The elapsed time from metaphase to chromatid scattering/cohesion fatigue was determined in HeLa-Slow cells (left) and HeLa-Fast cells (right) arrested at metaphase with ProTAME +/- 1.5μM STLC. Three independent experiments with a total of ≥150 cells were scored for each treatment and cell type. Error bars show standard deviation. The Mann-Whitney test was used for statistical analysis.

To reduce spindle tension we treated cells with low concentrations of the Eg5 inhibitor STLC (Skoufias et al., 2006). At high concentrations, STLC induces collapse of spindle poles. But, at reduced concentrations, spindles can be maintained with decreased interpolar distance and diminished spindle tension (Vallot et al., 2017). The decrease in spindle tension should reduce the outward force on kinetochores. In control cells, 1.5 uM STLC caused only a slight delay in normal mitotic progression (Supplementary figures 4D). In cells arrested at metaphase, STLC treatment led to significantly slower cohesion fatigue in both HeLa-Slow and HeLa-Fast cells (Figure C).

### Fatigued chromatids can congress to the metaphase plate

Normally metaphase in mitosis requires approximately 10 to 30 min before synchronous separation of sister chromatids in anaphase occurs, followed by mitotic exit. In contrast, when cells are experimentally delayed at metaphase, chromatids pull apart slowly and asynchronously while cells remain in mitosis (Figure 5A). The rate of chromatid separation varies widely among different cell lines. Chromosome spreads of cells arrested for a few to several hours (depending on the cell line) show complete separation of most chromosomes (Figure 1B). Typically, in cells that have undergone cohesion fatigue, some chromatids are oriented near the poles but many appear clustered near the metaphase plate (Figure 5A, last panel). To understand the behavior of chromatids during cohesion fatigue we used high resolution lattice light sheet microscopy to track chromatid movement after cohesion fatigue. Live imaging of LLC-Pk cells stably expressing GFP-Topoisomerase ||α, which marks both kinetochores and chromosome arms, revealed that partially and completely separated chromatids oscillate toward and away from the spindle midplane (Compare normal mitosis in Supplemental Video 1 and cohesion fatigue in Supplemental Video 2). Thus, unpaired kinetochores on chromatids separated by cohesion fatigue can subsequently align near the metaphase plate. This is likely due to formation of merotelic attachments of single kinetochores to microtubules from both poles and to microtubule-based ejection forces from the poles impacting chromatid arms.

**Figure 5:**
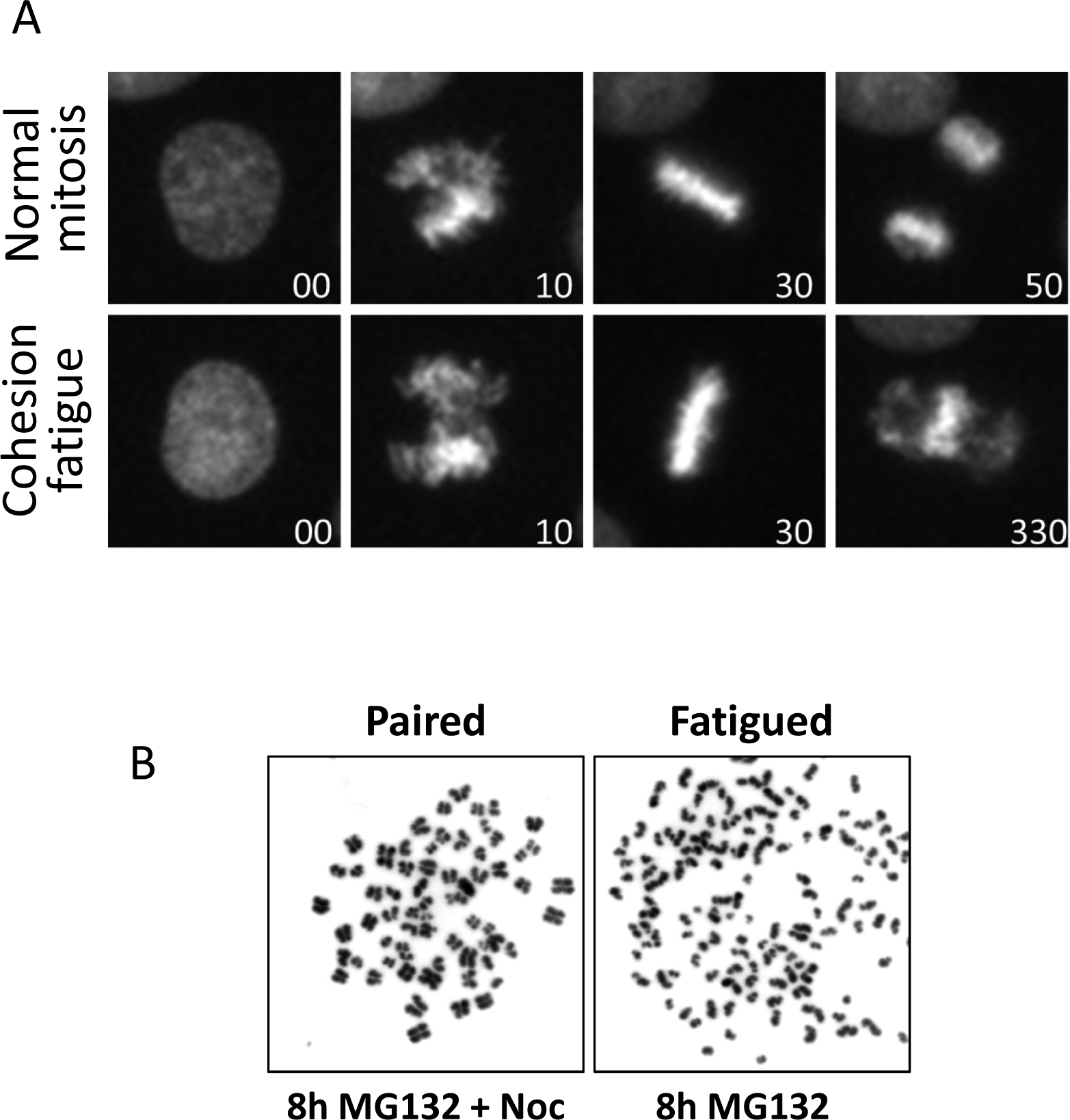
Separated chromatids can congress to the spindle midplane after cohesion fatigue. (**A)** Cohesion fatigue was assessed in HeLa cells stably expressing H2B-GFP treated with either DMSO (control) or ProTAME via live cell imaging. Time 00 indicates nuclear envelop breakdown (NEBD). DMSO-treated cells progress through normal mitosis (top panel), while ProTAME-treated cells undergo cohesion fatigue with a concentration of chromatin at the spindle midplane in the final image (bottom panel). **(B)** Chromosome spreads prepared from HeLa cells treated with MG132 ±Nocodazole for 8 h. Left panel shows paired sister chromatids (MG132+Nocodazole). Right panel shows separated sister chromatids (cohesion fatigue) after 8 h of metaphase arrest (MG132).

### Short delays at metaphase induce partial separation of chromatids at their kinetochores

Chromatid separation in cohesion fatigue is progressive, initiating at the kinetochores then advancing distally along the chromosome arms (Daum et al., 2011). To evaluate the time-course of chromatid separation after short delays, we tracked the interkinetochore distance between sister chromatids in LLC-PK cells. We detected significant separation of kinetochores in cells treated with MG132 for 3 h (Figure 6A) In most cells arrested for 3 h, sister chromatid arms remained attached, but kinetochores were significantly separated with many showing separations of more than 3 µm versus 1.75 µm in control metaphase cells and 0.7 µm in cells treated with 330 nM nocodazole (Figure 6A, 6B and Supplementary figures 5A). Like LLC-PK cells, HeLa-Slow cells also showed increased interkinetochore distances when delayed at metaphase for 3 h (Supplementary figure 5B). To examine the dynamics of chromatid separation in detail, we used live cell imaging to monitor metaphase-arrested LLC-PK cells expressing GFP-Topoisomerase IIα. As anticipated from the analysis of fixed cells, live cell tracking showed wider average distances between sister kinetochores in cells arrested at metaphase with MG132 for 3 hours compared to untreated control cells or cells treated for only 1 hour (Supplementary figures 5C). Moreover, cells treated with MG132 for 3 h showed a significantly larger range of stretching between sister kinetochores compared to cells treated for only 1 h. In metaphase of untreated cells or cells treated with MG132 for 1 h, the average distance between sister kinetochores varied over an average range of about 0.4 μm as sister kinetochores oscillated together and apart. In contrast, cells arrested at metaphase for 3 h showed a range of stretching between sister kinetochores of 1 μm or more (Figure 6C). Overall, moderate delays at metaphase cause abnormal separation of kinetochores.

**Figure 6:**
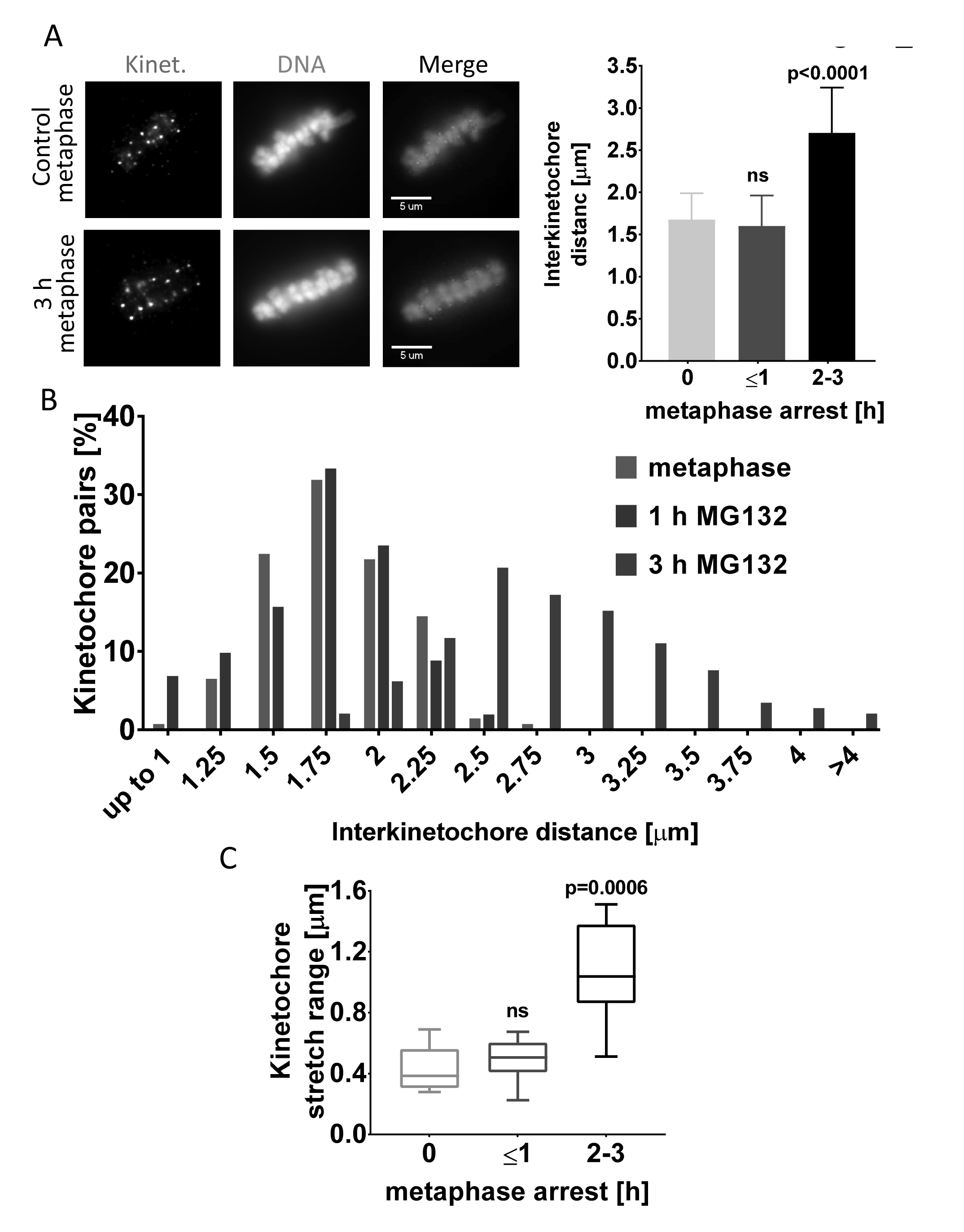
Sister kinetochores separate after transient metaphase arrest. **(A)** Transient metaphase arrest results in increased separation of sister kinetochores in LLC-PK cells. Representative immunofluorescence images (left) and quantification (right) of control metaphase cells or cells arrested at metaphase for up to 1 h and up to 2-3 h. The average distances between pairs of kinetochores from were compared in control metaphase cells and cells treated with MG132 for 1 or 3 h (n ≥ 125 kinetochore pairs in 5 cells from each treatment). One-way ANOVA, with Tukey’s multiple comparison test was used for statistical analysis. **(B**) The frequency distributions for distances between sister kinetochores from cells in (A) show increased proportions widely separated kinetochores in those arrested for 3 h. **(C)** The extent of stretching between sister kinetochores increases with time for cells arrested at metaphase. Live cell imaging determined the maximum stretching of sister kinetochores in LLC-PK cells arrested at metaphase for 1 or 2-3 h. For these measurements, n ≥ 10 pairs of kinetochores were imaged every 10 sec for 3 min. Kruskal-Wallis test with Dunn’s multiple comparison was used for statistical analysis.

### Partial separation of chromatids induces chromosome segregation defects

Transient delays in anaphase onset after most chromosomes have aligned at the metaphase plate often occur because one or more chromosomes lag in congression, even in an unperturbed, normal mitosis. To examine the immediate impact of partial chromatid separation that may occur during a transient delay, we arrested cells at metaphase, then released them into anaphase. We arrested LLC-PK cells with 5 µM MG132 for 3 h. Cells were washed into fresh medium without drug and then fixed 3.5 h later when most had entered anaphase. We examined every cell that entered anaphase for lagging chromosomes, anaphase bridges or micronuclei (Figure 7A left). Cells arrested at metaphase for 3 h with MG132 treatment exited mitosis with an error rate of ∼44%. Cells treated and released after a treatment with both MG132 and Nocodazole showed segregation errors in 18% of anaphases, significantly lower than MG132 treatment alone (Figure 7A right). Cells treated and released from a 3 h Nocodazole arrest exhibited a slightly elevated error rate of 7%. Untreated control cells exited mitosis with a missegregation rate of ∼4%. Because mitotic exit after release from MG132 requires approximately 3.5 h while recovery from Nocodazole takes only 30-60 min, cells released from the combination of MG132 and Nocodazole arrest at metaphase with an intact spindle for approximately 3 h. This finding is consistent with the higher rate of anaphase defects in these cells compared with controls. We also compared the accumulation of segregation defects in cells arrested at metaphase for different durations. We treated LLC-PK cells with MG132 for 1 or 4 h, released them in fresh medium and then evaluated the anaphases. In cells arrested for 1 h, 13% of the anaphases showed segregation errors, while in cells arrested for 4 h, 55% of revealed errors (Supplementary figures 6A).

**Figure 7:**
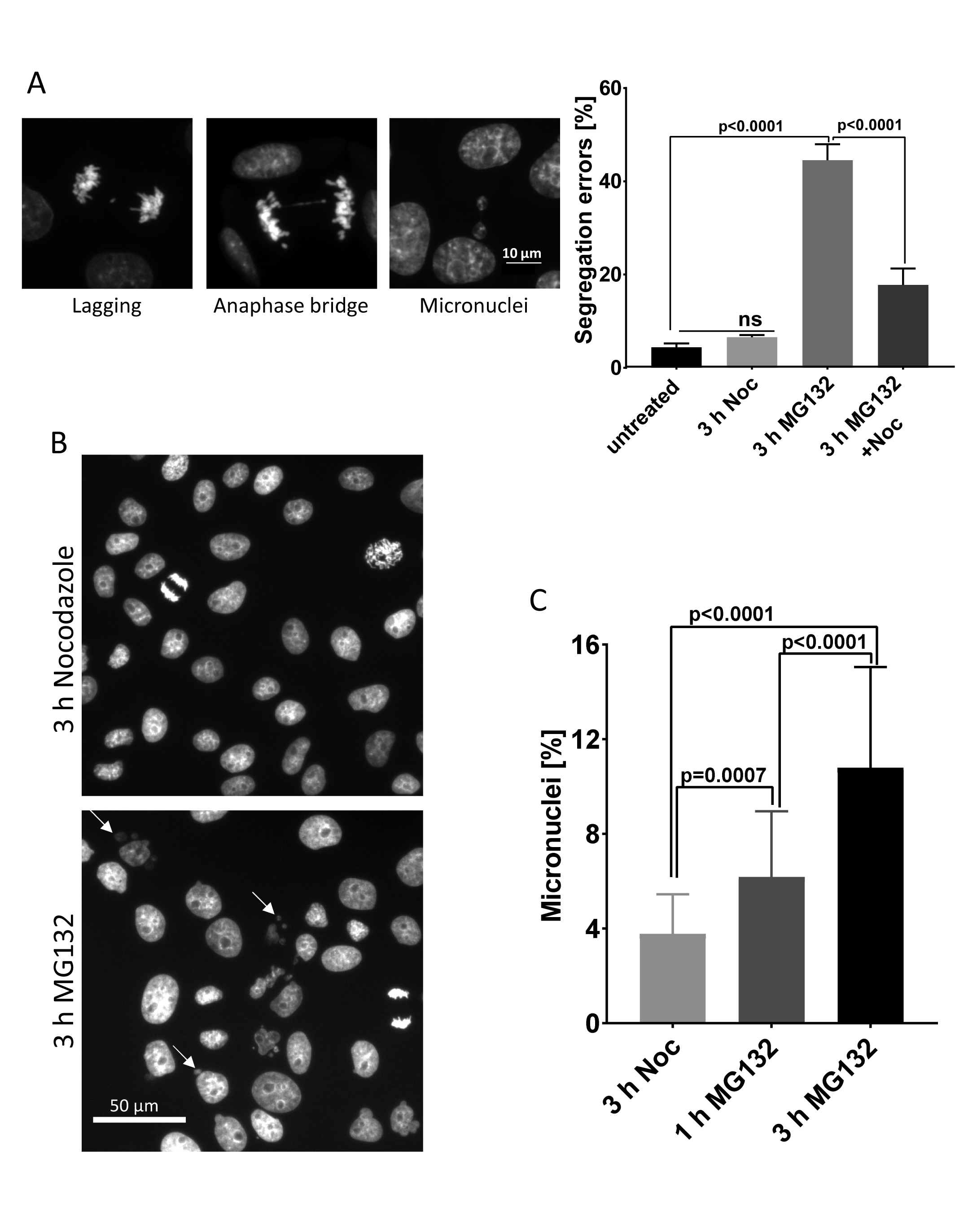
Transient metaphase delays induce segregation defects in LLC-PK cells. **(A)** Representative images (left) and quantification (right) of anaphase/telophase segregation defects (lagging chromosomes, anaphase bridges, or micronuclei) in LLC-PK cells transiently arrested at metaphase. Segregation defects during anaphase were examined in untreated cells or in cells transiently treated with Nocodazole, MG132, or MG132 +Nocodazole for 3 h in three independent experiments with >700 anaphases examined for each treatment. Error bars represent standard deviations. Ordinary one-way ANOVA with Holm-Sidak’s multiple comparison test was used for statistical analysis. **(B)** Transient delays at metaphase induce formation of persistent micronuclei. Low magnification images of LLC-PK cells transiently arrested at a prometaphase-like state with nocodazole or at metaphase with MG132 for 3 h then released for 24 h. Arrows indicate the micronuclei present in cells that were transiently delayed at metaphase. **(C)** Percentages of micronuclei in images from (B) were determined in ≥5000 cells from 50 randomly selected fields. One-way ANOVA was used for statistical analysis.

Not all anaphase chromosome segregation errors cause aneuploidy, as some lagging chromosomes are properly incorporated into daughter nuclei. However, missegregated chromosomes often decondense separately to form micronuclei that persist for long periods in daughter cells and can induce catastrophic DNA damage (Crasta et al., 2012; Hatch et al., 2013; Thompson and Compton, 2011; Zhang et al., 2015). We tested whether short metaphase delays increase the incidence of micronuclei. We quantified the number of micronuclei in LLC-PK cell cultures 24 h after transient arrests with MG132 or Nocodazole for 1 h and 3 h. Cells delayed at metaphase with MG132 treatment for 3 h exhibited significantly higher numbers of micronuclei compared to cells arrested for 1 h or cells treated with nocodazole for 3 h (Figures 7B and C).

To map the effects of short metaphase delays in greater detail, we used video microscopy. To achieve metaphase delays of varying lengths, we treated cells with 10, 20 or 30 µM ProTAME. ProTAME is a cell permeable inhibitor of the APC/C (Zeng et al., 2010). ProTAME delayed cells at metaphase in a dose-dependent manner for varying durations (Supplementary figures 6B). These ProTAME-induced metaphase delays were followed by three outcomes: 1) normal anaphase and mitotic exit, 2) defective anaphase involving lagging chromosomes, anaphase bridges, or micronuclei or 3) cohesion fatigue (Figure 8A). Cells delayed at metaphase for less than 2 h had a low incidence (14%) of anaphase defects. The number of cells showing defective anaphase increased to 37% in cells delayed for 3 h and to approximately 58% in cells delayed for 4 h. With longer arrest durations, the number of cells exhibiting defective anaphase declined, while the number of cells that underwent cohesion fatigue increased (Figures 8B and 8C). Cells exhibiting normal anaphase were delayed at metaphase an average of 134 ±111 min, while cells showing at least one kind of chromosome segregation defect were delayed for 199 ±106 min. Cohesion fatigue occurred in cells arrested at metaphase for 370 ±105 min (Supplementary figures 6C). Thus, cells showed an increased frequency of segregation errors that correlated with the duration of metaphase arrest but then exhibited cohesion fatigue after extended times at metaphase. Cells with massive chromatid separation did not generally enter anaphase, likely due to reactivation of the spindle checkpoint. Overall, limited separation of chromatids caused by transient metaphase delay produces chromosome segregation defects in anaphase and often generates micronuclei.

**Figure 8:**
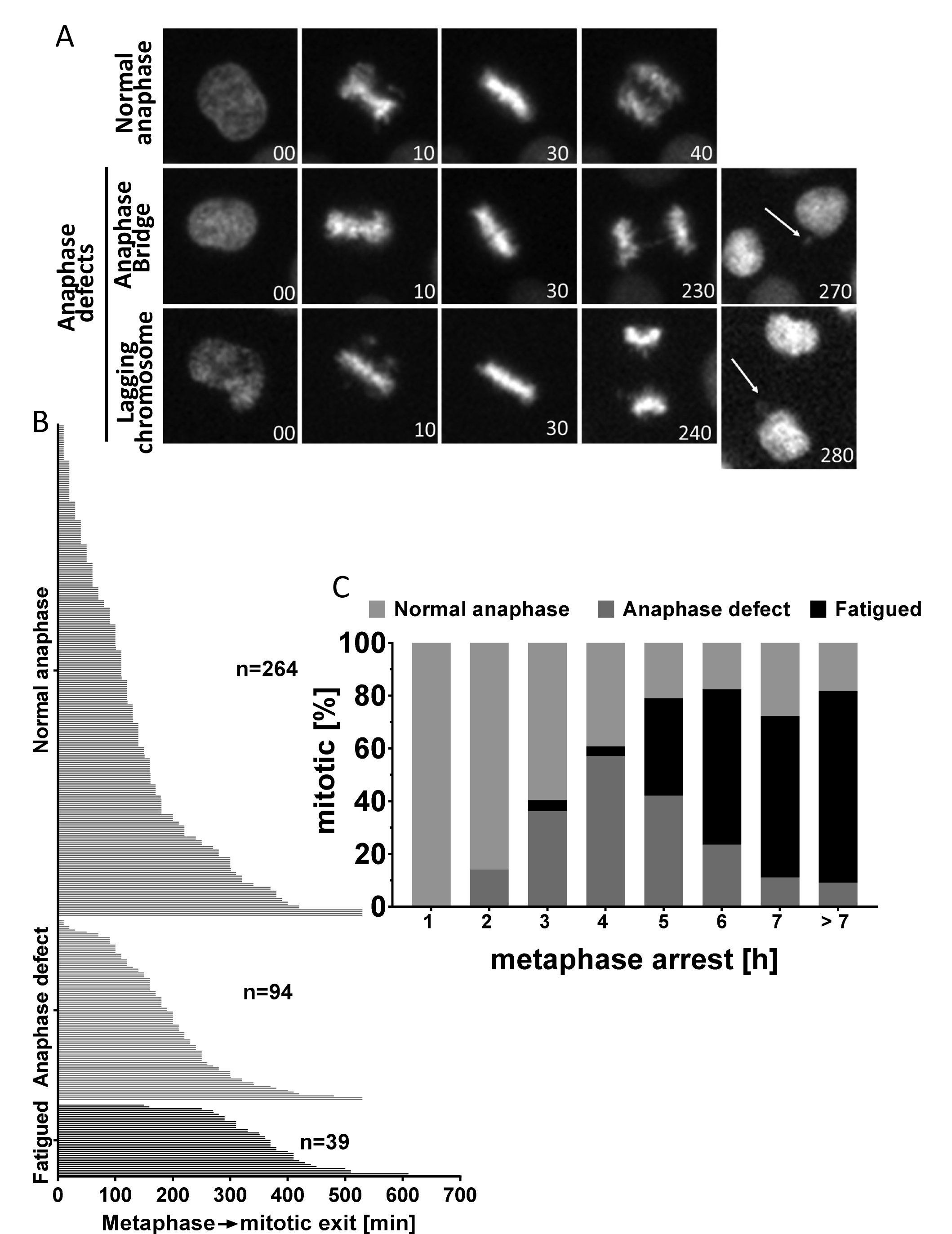
Segregation defects in HeLa cells scale with the length of metaphase delay. **(A)** Live-cell images of HeLa-H2B-GFP cells treated with 10, 20 or 30 μM ProTAME. Top panel shows a normal anaphase; bottom two panel show anaphase defects (arrows indicate micronuclei). **(B)** Fates of individual cells after ProTAME treatment. Most cells with slight delays had no segregation defects; intermediate delays increased the proportion of defective anaphase, while longer delays often resulted in cohesion fatigue. **(C)** Compiling of results for cells treated with ProTAME shows the increased anaphase defects at intermediate times of metaphase delay and increased cohesion fatigue at longer times.

## Discussion

Our data reveal that breaching of sister chromatid cohesion that accompanies cohesion fatigue does not require release of core Cohesin ring components from chromatids. It also does not appear to exploit a specific protein-protein interface in the Cohesin ring. More specifically, the Wapl-mediated opening of cohesin rings is not required after metaphase arrest to separate sister chromatids in cohesion fatigue. In contrast, loss of Wapl activity in early mitosis leads to increased retention of Cohesin on metaphase chromosomes, which does inhibit subsequent cohesion fatigue. Experimental manipulations that compromise Cohesin integrity in mitotic chromosomes accelerate cohesion fatigue. Our studies demonstrate the dynamic tension of the mitotic spindle, specifically the pulling forces acting on kinetochores is countered by the resistance of Cohesin that holds chromatids together. The rate of chromatid separation in cells delayed at metaphase yields a quantitative measure of these two antagonistic components. Our studies also reveal that partially or fully separated chromatids can travel to the spindle equator. This competence for the kinetochores of unpaired chromatids to congress to the metaphase plate was first described by (Brinkley et al., 1988). Finally, we show that in contrast to complete chromatid separation that accompanies cohesion fatigue, even relatively short metaphase delays can result in partial chromatid separation that lead to defects in chromosome segregation during the subsequent anaphase.

In mitosis in vertebrate cells, most Cohesin is released from chromosomes through the activities of mitotic kinases and the Wapl protein which act during early mitosis. Of the remaining Cohesin that remains bound to isolated mitotic chromosomes, most can be released by treatment with moderate levels of salt (Bermudez et al., 2012). Although not proven, we speculate that the functional or “cohesive” Cohesin that holds sister chromatids together reflects the minor, salt-resistant population. The precise molecular nature of Cohesin interactions with chromatin remains a topic of research and debate. For sister chromatid cohesion, most models are variations on two general modes of Cohesin/chromatin interaction termed “embrace” or “handcuff” models (reviewed in (Skibbens, 2016). Embrace models propose that both sisters are contained within the same Cohesin ring, while handcuff models suggest sister chromatids are enclosed in separate rings that are linked together. Recently a new model of chromatin binding by Cohesin was reported, termed “hold-and-release” (Xu et al., 2018). The hold-and-release model proposes that DNA is sandwiched by arched coiled-coils of SMC components rather than entrapped within a ring. Although sister chromatids undergo separation during cohesion fatigue, we found no change in the amount of salt-stable, core Cohesin components bound to chromosomes before or after chromatid separation (Figure 1). This surprising result suggests that salt-resistant Cohesin is not released from chromatids during their separation. This outcome is most consistent with the handcuff models or the newly suggested hold-and-release model of interchromatid cohesion since these do not necessarily require Cohesin release at chromatid separation. However, there are other potential explanations for our findings. One is that sister chromatid cohesion is mediated only by a minor subfraction of the salt-resistant Cohesin, which is indeed released during cohesion fatigue at levels we cannot detect. Recent work in Drosophila, where total Cohesin levels were genetically regulated, shows that expression of very low levels of Cohesin can maintain normal sister chromatid cohesion at metaphase (Oliviera, personal communication). Another possibility to explain our results is that Cohesin interactions with chromatin are remodeled during fatigue from binding sister chromatids together to binding chromatin segments within a single chromatid. This possibility has been proposed to explain centromere structure in budding yeast (Yeh et al., 2008) and chromatin immunoprecipitation studies in yeast suggest that Cohesin may be able to break and reform chromatid linkages during mitosis (Ocampo-Hafalla et al., 2007). Alternatively, salt-resistant Cohesin might be released during chromatid separation only to rebind onto the separated chromatids later. Finally we cannot exclude the possibility that DNA damage generated during cohesion fatigue could recruit Cohesin during separation (Unal et al., 2004).

Another potential explanation for the retention of Cohesin on chromatids during cohesion fatigue is that protein-protein interactions among proteins comprising the Cohesin ring may open transiently. Opposing poleward-directed tension may exploit these transient openings, allowing sister chromatids to slip apart. Separation would initiate at the kinetochores then progress down the length of the chromosome arms, the exact behavior observed in time lapse imaging (Daum et al., 2011). However, in our current study, after allowing the normal activity of the Wapl-mediated Cohesin release in prometaphase, locking of individual gates of Cohesin complexes at metaphase did not inhibit cohesion fatigue (Figure 2E). This evidence suggests that neither the SMC3-RAD21, Wapl gate nor the SMC1-SMC3 and the SMC1-RAD21 gates are individually required for cohesion fatigue. Because we cannot lock multiple gates at the same time, we cannot eliminate the possibility that transient openings of multiple Cohesin gates contributes to cohesion fatigue. In our previous study we showed, by siRNA depletion, that background Separase activity did slightly affect the timing of cohesion fatigue but was not required (Daum et al., 2011). This evidence, along with our current Aurora B inhibition studies (Figure 2C) and gate locking experiments suggest that the mechanism of cohesion fatigue is likely a novel, yet undiscovered pathway of breaching sister chromatid cohesion. Thus, the phenomenon of cohesion fatigue provides an additional tool to study the nature of Cohesin interaction with chromatin.

Our studies reveal that the amount of Cohesin retained on chromosomes dictates the rate of cohesion fatigue. While Wapl activity is not required for cohesion fatigue, previous work showed that siRNA-mediated depletion of Wapl decreased the rate and extent of cohesion fatigue (Daum et al., 2011; Stevens et al., 2011). We tested the idea that this was due to increased retention of Cohesin on mitotic chromosomes by using HeLa cells stably expressing SMC1-GFP and showed directly that cohesion fatigue is delayed in cells where mitotic chromosomes show enrichment of Cohesin after Wapl depletion prior to mitotic entry. We also showed that expression of the Wapl antagonist, Sororin, mutated to be resistant to removal by mitotic phosphorylation, also delays cohesion fatigue. These experiments confirm that inhibition of Wapl-mediated prophase pathway before mitotic entry enriches Cohesin on chromosomes and delays cohesion fatigue (Figures 2A and 2B).

While our results show that the Wapl-mediated cohesin removal is not essential for cohesion fatigue, it may still influence timing. Complete disruption of spindle microtubules eliminates cohesion fatigue. However, extended arrest in the absence of microtubules renders cells more sensitive to subsequent cohesion fatigue when metaphase spindles are allowed to form (Figure 3A). Over time in cells arrested with depolymerized spindles, chromosome-associated Cohesin may diminish through continued Wapl-mediated Cohesin release and/or background separase activity. In support of this idea, we found diminished Cohesin levels on chromosomes isolated from cells treated with Nocodazole for longer times (Supplementary figures 3C).

Two previous studies reported that experimentally induced metaphase delays would reduce immediate chromosome segregation errors (Cimini et al., 2003; Ertych et al., 2014). However, our detailed analyses indicated the opposite, that transient metaphase delays increase the incidence of segregation defects during anaphase (Figures 7 and 8). The reason for this discrepancy is not clear, but the previous reports used different cell lines and different experimental conditions. Consistent with the previous studies, we found that very short metaphase delays did not increase the chromosome segregation errors (Supplementary figures 6C). Our results point to a critical threshold of metaphase arrest and accompanying kinetochore separation that may be required before the delay becomes detrimental. In our experiments increased kinetochore separation abrogates the normal back-to-back orientation of sister kinetochores allowing greater chances for merotelic attachments of single kinetochores to both spindle poles. Such merotelic kinetochore attachments have been shown to increase the incidence of anaphase defects (Cimini et al., 2003; Salmon et al., 2005; Thompson and Compton, 2011). Cohesion fatigue that generates chromatid separation in several chromosomes likely compromises cell viability either through cell cycle arrest mediated by the spindle checkpoint or through catastrophic chromosome missegregation if cells exit mitosis. The more subtle errors that accompany partial chromatid separation for cells with shorter delays at metaphase may produce segregation defects that can propagate in daughter cells.

Although certain cells, notably some cancer cell lines, exhibit high degrees of chromosome instability, most dividing cells in culture show low rates of spontaneous segregation errors that can manifest in several ways (Thompson and Compton, 2008). As with the other errors, we observe spontaneous cohesion fatigue in normal control cells at low frequency (data not shown). Cells that undergo substantial cohesion fatigue with many separated chromatids are likely to die, because separated single chromatids elicit spindle checkpoint signaling (Lara-Gonzalez and Taylor, 2012), which promotes continued mitotic arrest and cell death. Even if they survive and divide after delay, the progeny cells would have highly abnormal chromosome content and would likely be inviable. In this study we also focused on less extreme circumstances, where shorter delays at metaphase allowed sister chromatids to partially separate before anaphase onset (Figure 6).

Cohesin complexes contain one of two auxiliary “Stromal Antigen” components, SA1 or SA2. In HCT116 and RPE1 cells, depletion of SA2 causes significant increases in lagging chromosomes (Kleyman et al., 2014). Furthermore, in HCT116 cells, knockout of the Stag2 gene, which codes for SA2, does not strongly affect normal mitotic progression but may increase the incidence of aneuploidy (Solomon et al., 2011). We found that HCT116 cells lacking SA2 underwent faster cohesion fatigue compared to parental HCT116 cells (Figure 3B). However, it does not appear that release of SA2 accompanies cohesion fatigue in normal cells, since chromatin-associated SA2 does not decrease during metaphase arrest or after chromatid separation (Figure 3C). Inactivating mutations in the Stag2 gene are correlated with aneuploidy in some cancers (Solomon et al., 2011). We propose that an increased propensity for full or partial chromatid separation due to cohesion fatigue may contribute to aneuploidy in cells with mutations in Stag2.

Previous studies suggested that cells prone to undergo rapid cohesion fatigue showed altered distribution and reduced levels of the Cohesin protector protein, SGO1 (Tanno et al., 2015). However, both fast and slow fatiguing HeLa cells exhibited no significant reduction in SGO1 protein during cohesion fatigue when compared to metaphase levels (Supplementary figures 1D and 1E). SGO1 levels were higher at metaphase in HeLa-Fast cells compared to HeLa-Slow cells, the opposite one might expect if Sgo1 levels regulated the rate of cohesion fatigue. Taken together, our observations indicate that breaching cohesion during cohesion fatigue may not require canonical Cohesin removal mechanisms, such as the Wapl pathway or the activity of Separase. However, we believe it is highly likely that these mechanisms may influence the sensitivity and rates of cohesion fatigue in different cells and under different conditions.

Metaphase is a point of balance between microtubule-dependent pulling forces that separate chromatids versus cohesive forces that hold chromosomes together. We show that mitotic spindle microtubule dynamics affect cohesion fatigue, likely by modulating spindle-pulling forces. Low concentrations of Taxol accelerate fatigue, while low concentrations of Nocodazole slow it. Partial inhibition of Eg5 kinesin with STLC likely compromises overall spindle tension to relax spindle-pulling force and slow cohesion fatigue (Figure 4). These results highlight the roles of robust and dynamic microtubules and sufficient spindle-pulling forces to separate sister chromatids during cohesion fatigue.

While normally transient, metaphase can be delayed. Our data suggest that many factors contribute to cell sensitivity to cohesion fatigue including the various canonical Cohesin regulators. However, a complete understanding of the primary molecular mechanisms underlying chromatid separation during cohesion fatigue remains unresolved and may reflect an incomplete understanding of sister chromatid cohesion. Future studies of cohesion fatigue may provide insight into the nature of Cohesin complex-chromatin interactions. While complete chromatid separation of many chromosomes will likely result in cell death or inviable progeny cells, complete separation of one or a few chromosomes and/or partial chromatid separation may be an important source of genomic instability that perpetuates the evolution of malignant cells in cancer.

## Material and methods

### Cell culture and drug treatments

HeLa, LLC-PK, HCT116 and HEK293 cells were cultured in flasks in DMEM-based media supplemented with 2 mM HEPES, non-essential amino acids (NEAA), sodium-pyruvate, 1X penicillin-streptomycin (P/S, Corning, 30-002-CI) and 10% FBS. Cells were maintained at 37°C in 5% CO_2_ in a water-jacketed incubator. Cells were subcultured every other day and were used within 6 months of thawing from liquid nitrogen. Unless otherwise specified, drugs were applied at the following concentrations: Nocodazole: 330 nM, MG132: 25µM, ProTAME: 25 µM, Rapamycin: 100 nM, ZM447439: 2.5 uM. All cell lines were routinely tested for mycoplasma. HeLa cells, LLC-PK cells and HCT116 cells were mycoplasma free. The HEK293 cells were found to be mycoplasma positive. Unfortunately, all the stock cultures, even the earliest isolates at the laboratory of origin were found to be mycoplasma positive. We used several approaches designed to cure the mycoplasma contamination, but these were unsuccessful.

### Chromosome/chromatin isolation

Subconfluent cultures of HeLa cells were treated with Nocodazole for 12 - 16 h, then mitotic cells were collected by shake-off. Cells remaining in the flasks (interphase cells primarily in G2) were collected by trypsinization. Cells were centrifuged in 50 ml tubes at 200 Xg for 4 min and re-suspended in warm media at 1X10^6^ cells/ml. 4×10^5^ cells were aliquoted into 1.5 ml micro-centrifuge tubes and centrifuged at 200 Xg for 5 min. The cell pellet was then lysed with cold Extraction Lysis Buffer (ELB) by repeated pipetting. The ELB contained PHEM buffer: 60 mM <u>P</u>IPES, 25 mM <u>H</u>EPES, 10 mM <u>E</u>GTA and 4 mM <u>M</u>gCl_2_ with 0.1 M NaCl, 1% CHAPS, 1mM DTT, and 1:200 protease inhibitor (Sigma, P8340). Lysed cells were incubated for 20 min in ice then centrifuged at 1400 Xg for 10 min. A fraction of soluble supernatant was saved. The pellets were subjected to two more cycles of resuspension in ELB and centrifugation.

### Western blot

Supernatant and chromatin pellets were dissolved in 1X loading buffer (1X LDS sample buffer (Thermofisher, NP007) + 50 mM DTT). Equivalent cell numbers were loaded on 4-12% NuPAGE gels, electrophoresed at 50 V for 7 min, then for 2 h at 150 V in MOPS SDS running buffer. Proteins were transferred onto 0.45 micron PVDF membrane in transfer buffer (50 mM Tris, 192 mM Glycine and 0.05% SDS) containing 15% methanol with a Midi transfer apparatus (Idea Scientific). Blots were blocked with 5 % non-fat dry milk in PBST (PBS with 0.05% Tween 20) or 1:10 Sea Block (Thermofisher, 37527). Blots were cut into pieces and incubated with rabbit anti-SMC3 (Bethyl, A300-055A) at 1:1000 in block, rabbit anti-RAD21 (Bethyl, A300-080A, BL331) at 1:1000, mouse anti-SA2 (Santa Cruz, J-12) at 1:1000, rabbit anti-CENPA (Millipore, 07-574) at 1:200 and rabbit anti-Histone H3 (Abcam, ab1791) at 1:10000 at 4°C overnight with gentle rocking. Blots were washed 3 times with PBST then labeled with Horseradish Peroxidase (HRP) conjugated goat anti-rabbit secondary (JacksonImmunoresearch, 11-035-144) at 1:20000. For far red fluorescent detection, goat anti-rabbit, or anti-mouse (Azure biosystem, AC2128 and AC2129) were used at 1:10000 at room temperature for 2.5 h. Blots were washed again 3 times with PBST. For HRP detection blots were treated with Pierce West Pico reagent for 5 mins, then captured by chemiluminescence with a Kodak 4000R Image Station. For far-red fluorescence, membranes were imaged using an Azure c600 imaging system. Blot quantification was done using the raw images with Metamorph Software (Molecular devices).

### Chromosome spreads

Mitotic cells were washed with warm media by centrifuging at 300Xg for 3 min. Cells were suspended in 500 µl of warmed swelling buffer (40 % complete media + 60% DI water). Samples were incubated in a 37°C water bath for 15-18 min. Swollen cells were fixed by adding 1 ml 3:1 methanol: acetic acid, then incubated for 10 min. The cells were pelleted for 5 min at 250Xg, then washed with 1 ml fixative and pelleted once more. The cell pellets were resuspended in 100-200 µl fixative, then 40-50 µl of cell suspension was dropped from a height of 60 cm onto a 22 mm^2^ coverslip that was cleaned with 95% ethanol and wiped with acetic acid. The coverslips were immediately placed inside a 150 mm plastic culture dish on top of wet filter paper. The lid was left off, and the coverslips were allowed to dry in the humidified chamber. Once dried, coverslips were stained with DAPI (100 ng/ml) and SYBERGold nucleic acid dye (1:20000). Slides were imaged with a Zeiss Axioplan II microscope using a 100X objective, Hamamatsu Orca II camera and Metamorph software. At least 200 mitotic spreads were scored for each sample. If an individual cell spread had more than 10 single chromatids, the cell was scored as fatigued.

### Live cell imaging

Cells were grown in chambered cover glasses (Lab-Tek) for 24 h, then the medium was changed to L-15 phenol red-free medium supplemented with P/S, NEAA and 10% FBS. The surface of the medium was overlayered with mineral oil to reduce evaporation. For most experiments, chambers were transferred to a Zeiss Axiovert microscope then imaged while using an air-curtain heater to maintain the temperature at 37°C. Images were acquired every 7-10 min for 18-20 h with a Zeiss 20X objective and ORCA-ER Hamamatsu camera using Metamorph Software (Molecular Devices LLC). Images were analyzed using Metamorph software. For experiments; Wapl depletion in SMC1-GFP cells, STLC treatment in HeLa-Fast, Taxol treatment in HeLa-Slow, Rapamycin treatment in HeK293, ZM447439 treatment, and Sororin mutant images were acquired using 20X objective in a Nikon Ti microscope fitted with an OKOlab environmental chamber. For each cell that entered mitosis, the intervals from nuclear envelope breakdown (NEBD) to metaphase and to anaphase onset or cohesion fatigue were recorded. To induce metaphase arrest, cells were treated with 25 µM MG132 or 25 µM ProTAME, and scored as fatigued when approximately 10% of the chromosomes had undergone chromatid separation.

For high resolution cohesion fatigue imaging, LLC-PK cells constitutively expressing EGFP-Topoisomerase II alpha were grown on round 5mm glass coverslips in DMEM-based media to densities between 60% and 80%. For control cultures exhibiting normal mitotic progression DMSO was added to culture medium at 0.1%. To induce cohesion fatigue the 26S proteasome inhibitor, MG132, dissolved in DMSO was added at 10 uM to experimental cultures which resulted in a 0.1% DMSO concentration. Prior to image acquisition in order to improve fluorescence capture and to remove the requirement for carbon dioxide pH buffering, media was exchanged to phenol-free Leibovitz L-15 media with L-glutamine (Cat.# AT207-1L, VWR), 10 % FBS, Penicillin, Streptomycin with or without MG132 as described above. Fluorescence images of EGFP-Topoisomerase II alpha at 5 second intervals encompassing the entire cell volume were acquired using the lattice light sheet microscope (LLSM) at Janelia Research Campus’s Advanced Imaging Center. Movies were prepared using Imaris software.

### siRNA experiments

HeLa cells stably expressing SMC1-GFP were grown on chambered cover glasses. Cells were transfected with Wapl siRNAs #1 GAGAGAUGUUUACGAGUUU; #2 CAACAGUGAAUCGAGUAA or universal negative control (Sigma catalog # SIC001) using RNAi Max lipofectamine (Thermofisher catalog # 13778150). 48 h after transfection, live cell imaging was done as described above. Only cells that showed a clear GFP signal on metaphase chromosomes were included indicating significant depletion of Wapl. The elapsed time from NEBD to metaphase and to chromatid separation was measured.

### Immunofluorescence

Cells grown on 22 mm^2^ coverslips were simultaneously fixed and permeablized with 2% PFA and 0.5% Triton X-100 in 1X PHEM buffer at room temperature for 15 min. The cells were blocked with 20% boiled normal goat serum (BNGS) for at least 20 min. Coverslips were incubated with primary antibody, CREST (1:800, Antibody INC, 15-134), rabbit anti SGO1 (1:500, a gift from Dr. Hongtao Yu) diluted in 5% BNGS in PBST overnight at 4°C. Coverslips were washed 3 times with MBST (MOPS buffered saline with 0.05% Tween 20), then incubated in secondary antibody, goat anti-rabbit conjugated to CY3 at 1:1500 (JacksonImmuno, 111-165-045109), goat anti-human conjugated to FITC at 1:800 (JacksonImmuno, 109-95-088) for 2 h at room temperature. After incubation with secondary antibodies, coverslips were washed three times again then labeled with DAPI (100 ng/ml) for one min. Coverslips were mounted on slides with Vectashield mounting media (Victor labs, H-1000), then sealed with clear nail polish. Fluorescence images of cells were taken using a Zeiss Axioplan II microscope with a Zeiss 100X objective, Hamamatsu Orca II camera and Metamorph software. Distances between pairs of kinetochores were measured using the region measurement tool in Metamorph software.

### Transient metaphase arrest

For fixed cell analysis, LLC-PK cells grown on 22mm^2^ were treated with 5 µM MG132 ± 330 nM Nocodazole for 3 h. Arrested cells were washed 4 times with warm DMEM then released into complete DMEM medium to complete mitosis. 3.5 h after release from drug, cells were fixed with 3:1 methanol: acetic acid and labelled with (DAPI 100 ng/ml). Anaphase cells were examined visually for lagging chromosomes or anaphase bridges with a Zeiss Axioplan II and a Zeiss 100X objective. For identification of micronuclei, LLC-PK cells grown on coverslips were transiently arrested with Nocodazole for 3 h or MG132 for 1 or 3 h then washed and released into complete medium. 24 h after release, cells were fixed with 3:1 methanol: acetic acid then labeled with DAPI (100 ng/ml). Each coverslip was imaged at 50 random positions with a Zeiss Axiovert microscope and Zeiss 20X objective. The total number of cells and micronuclei in a field was quantified using Metamorph software. For live cell imaging, HeLa-H2B-GFP cells grown on chambered cover glasses were treated with 10, 20, or 30 µM ProTAME in L-15 medium then imaged every 10 min for 18 h. Every cell that entered the mitosis was examined visually at anaphase for any visible signs of anaphase bridges, lagging chromosomes or micronuclei formation.

## Acknowledgements

We thank Dr. Arshad Desai for HeLa cells expressing H2B-mRFP, Dr. Kyoko Yokomori for HeLa cells expressing SMC1-GFP, Dr. Todd Waldman for HCT116 SA2 knockout cells, Dr. Olaf Stemmann for HeK293 cells expressing FRB- and FKBP-fused Cohesin proteins, and Dr. Hongtao Yu for HeLa cell expressing 9A-mutnat Sororin and anti-SGO1 antibodies. We also thank John Heddleston and Teng-Leong Chew at Janelia Research Campus’s Advanced Imaging Center and HHMI and More Foundation for providing instrumentation and technical assistance for lattice light sheet microscopy imaging. Our thanks to all the members of Program in Cell Cycle and Cancer Biology at OMRF for insightful discussion and comments. This work was supported by the National Institute of General Medical Sciences (R01GM111731) and by the McCasland Foundation.

## Supplemental Figure Legends

**Supplemental figure 1: Buffers containing moderate levels of salt removes most Cohesin from isolated mitotic chromosomes, and Sgo1 levels do not correlate with cohesion fatigue. (A)** Titration of Cohesin SMC3 and CENP-A used for quantification of Cohesin proteins. Dilutions of the 0 h sample were blotted with samples from other treatments and time points. The linear ranges of dilutions were determined for accurate quantification of the Cohesin component SMC3 (top) and loading control CENP-A (bottom). The vertical dotted lines represent the number of cells that generate linear protein loaded to signal ratios. 10000 cells equivalent protein was loaded for subsequent experiments. **(B)** Immunoblot of Cohesin components (SMC3 and RAD21) from chromosome/chromatin fractions of mitotic and interphase (G2) HeLa cells isolated in buffers with increasing NaCl. Cohesin was readily released from mitotic chromosomes compared to interphase chromatin by salt treatment. More than 90% of Cohesin was released from mitotic chromosomes by treatment with 100 mM NaCl. **(C)** Total SGO1 levels relative to CREST signals were determined by immunofluorescence in HeLa-Slow and HeLa-Fast cells. At least 10 cells for each cell line were quantified. **(D)** SGO1 levels determined by immunofluorescence in HeLa-Slow cells that were treated with MG132 for 6h. **(E)** Total SGO1 levels in HeLa-Fast cells treated with MG132 for 3h. For each cell line and treatment, at least 5 randomly selected cells were imaged with 0.5 μm Z sections. Summed projections were generated with Metamorph Software using the stack arithmetic tool. Region of DAPI staining was used as the mask to quantify total chromosome-associated SGO1.

**Supplemental figure 2: Inhibition of the prophase pathway before mitotic entry enriches chromosome-bound Cohesin and delays cohesion fatigue. (A)** Images from live cell imaging of HeLa-SMC1-GFP cells depleted of Wapl by RNAi. Wapl-depletion resulted in cells showing strong SMC1-GFP signals on mitotic chromosomes. **(B)** The elapsed times from NEBD to metaphase were determined in Hells treated with 0.5 uM ZM447439. ZM Treatment caused significant delays in chromosome alignment indicating Aurora B inhibition. >100 cells for each treatment were scored. **(C)** Chromosome spread on the left shows unresolved chromosome arms in a cell where the Wapl-mediated cohesin release was blocked by locking SMC3-RAD21 interface prior to entrance of cells into mitosis. Spread on right is from a control cell of the same type without Rapamycin addition (unlocked SMC3-RAD21 gate). **(D)** Percentages of chromosome spreads with closed/unresolved chromosome arms in HEK293 cells expressing pairs of Cohesin fusions to which Rapamycin was added to lock different Cohesin gates. Unresolved arms were increased only when the SMC-RAD21 gate was locked. **(E)** Elapsed times form metaphase to cohesion fatigue were determined in three gate-locking Hek293 cells line with gates locked prior to mitotic entry. Locking the SMC3-RAD21 gate but not other gates delayed cohesion fatigue.

**Supplemental figure 3: Longer mitotic arrest reduces the levels of chromosome Cohesin and the amount of chromosome-associated Cohesin is reduced in cells lacing SA2. (A)** Longer mitotic arrest in Nocodazole results in greater separation of chromosome arms. Cells were arrested with Nocodazole for 3 h (left panel) or 14 h (right panel) then prepared for chromosome spreads. **(B)** Cells that take longer to align their chromosomes fatigue faster. Elapsed times from metaphase to cohesion fatigue in STLC+ProTAME-treated cells plotted against times from NEBD to metaphase. **(C)** A small reduction of salt-resistant SMC3 in chromosome fractions accompanies longer arrest in Nocodazole. Immunoblot of chromosome fractions from mitotic HeLa cells arrested in mitosis with Nocodazole for 5h or 11h. Graph shows quantification. **(D)** SA2 knockout HCT116 cells lack expression of detectable SA2 protein. Immunoblot with anti-SA2 antibody of whole cell lysates of SA2 knockout HCT116 cells and parental HCT116 cells. **(E)** Core Cohesin ring protein levels are reduced in SA2 knockout cells relative to parental cells. Immunoblot of salt-treated chromosome fractions from HCT116 SA2 knockout and parental cells. Graph shows band intensity of SMC3 and RAD21 relative to CENP-A.

**Supplemental figure 4: HeLa-Fast and HeLa–Slow cells differ in fatigue kinetics, and HeLa-Slow cells undergo slower mitosis in low concentrations of spindle drugs. (A)** The elapsed time from metaphase to chromatid separation (cohesion fatigue) was determined from live cell imaging in HeLa-Slow and HeLa-Fast cells treated with 25 uM ProTAME. Average times of fatigue with standard deviations are shown from three independent experiments with ≥300 cells. Mann-Whitney test was used for statistical analysis. **(B)** Kinetics of cohesion fatigue in cells from (A). **(C)** HeLa-Slow cells treated with low concentrations of Nocodazole (5 nM) or Taxol (1.5 nM) show slowed progression through mitosis. The elapsed times from NEBD to anaphase was determined by live cell imaging. **(D)** Low concentration of STLC slowed but did not block mitotic progression in HeLa-Slow cells. The elapsed time from NEBD to anaphase was determined with cells treated with 1.5 μM STLC with ≥100 cells per treatment scored.

**Supplemental figure 5: Short metaphase arrest causes increased interkinetochore distance only in the presence of a functional spindle. (A)** Interkinetochore distances were measured in LLC-PK cells treated with MG132 and Nocodazole for 1 or 3 h compared to normal metaphase (Control). A total of ≥100 kinetochore pairs from 5 cells were measured. One-way ANOVA was used for statistical analysis. **(B)** Interkinetochore distances were measured in HeLa cells arrested at metaphase with MG132 for 30 min or 3 h. A total of ≥100 kinetochore pairs from 5 cells were measured. **(C)** Interkinetochore distances were determined from live cell imaging in LLC-PK cells stably expressing Topo-II-GFP. Cells were arrested in metaphase with MG132 treatment for 1 or 3 h. Transient metaphase arrest led to significant kinetochore separation if spindle microtubules were intact.

**Supplemental figure 6: MG132 and ProTAME induce metaphase delays whose duration determines cell fate. (A)** Percentages of defective anaphases in LLC-PK cells treated with MG132 for 1 or 4 h then released into drug-free medium. **(B)** Metaphase arrest durations were determined from live cell imaging of HeLa-Slow cells treated with increasing concentrations of ProTAME. For each concentration of ProTAME, ≥75 cells were scored. The Kruskal-Wallis test was used for statistical analysis. **(C)** The average length of metaphase arrest was determined in cells from (B) that exited mitosis to 1) normal anaphase, 2) defective anaphase, or 3) cohesion fatigue.

## Supplemental Videos

**Supplemental Video 1:** Normal mitotic progression visualized by lattice light sheet microscopy in LLC-PK cells stably expressing EGFP-Topoisomerase IIα, which highlights kinetochores and chromosome arms. Time = min:sec.

**Supplemental Video 2:** Consequences of cohesion fatigue visualized by lattice light sheet microscopy in LLC-PK cells stably expressing EGFP-Topoisomerase IIα. Imaging was initiated 6 h after metaphase arrest induced by treatment with MG132. Some chromatids (00 min, middle left) have separated completely. Other chromosomes show extensive separation of their kinetochores but retain partially cohered arms. Separated kinetochores undergo oscillations toward and away from the spindle equator. Time = min:sec

