## Supplementary figures and images for "Multiple Determinants and Consequences of Cohesion Fatigue in Mammalian Cells" (from another version of this preprint: version 2, 2018-03-27)

### Supplementary Materials

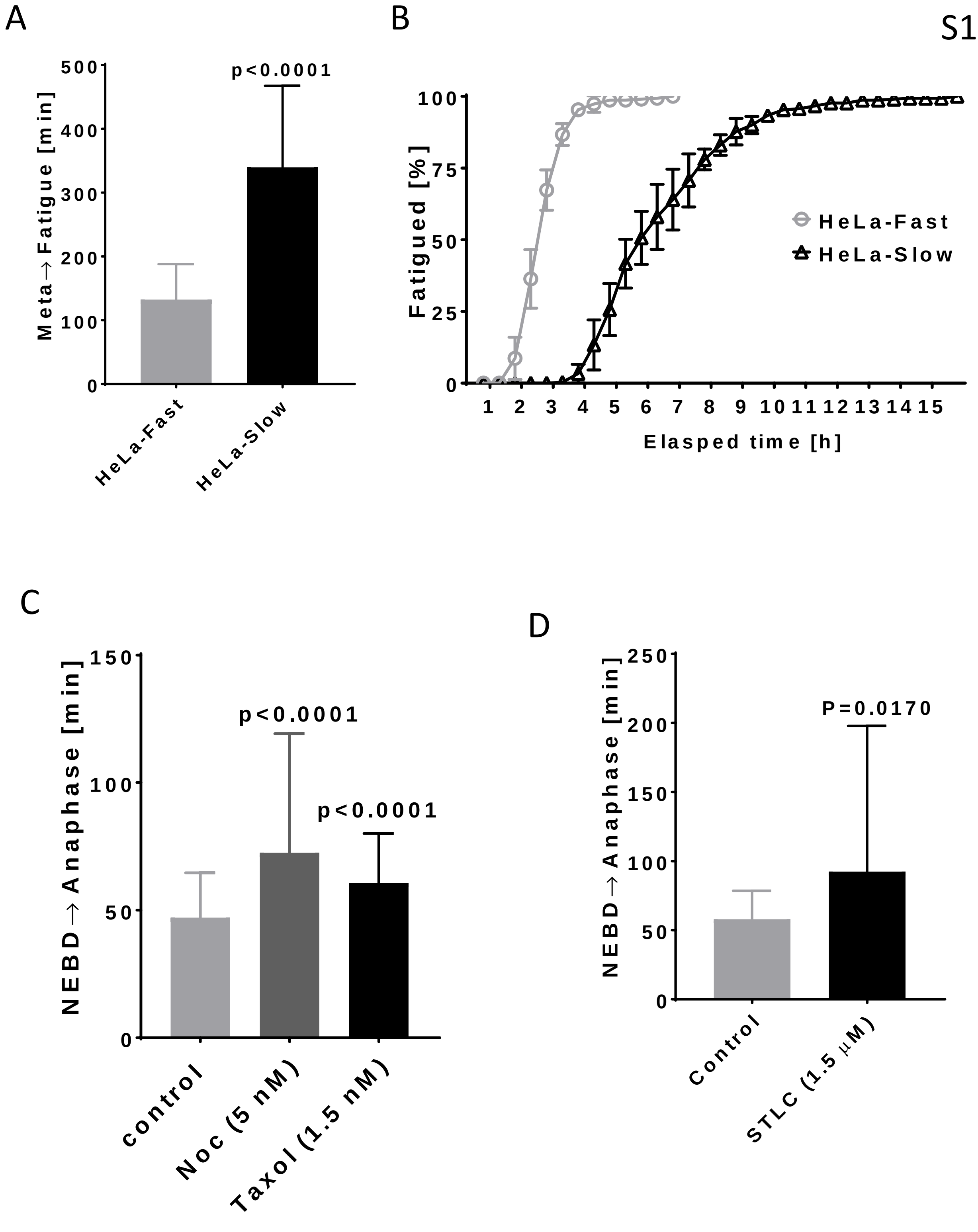

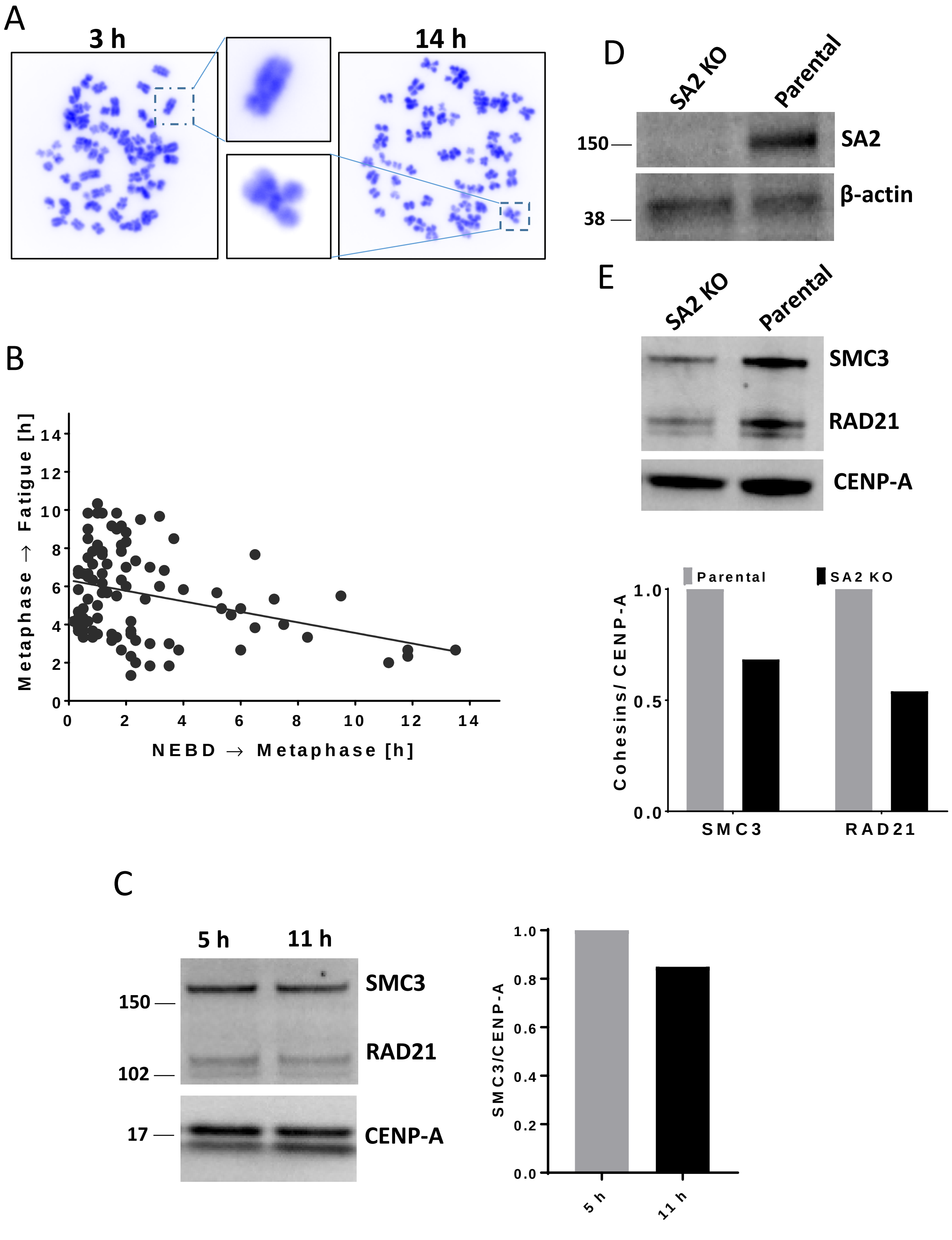

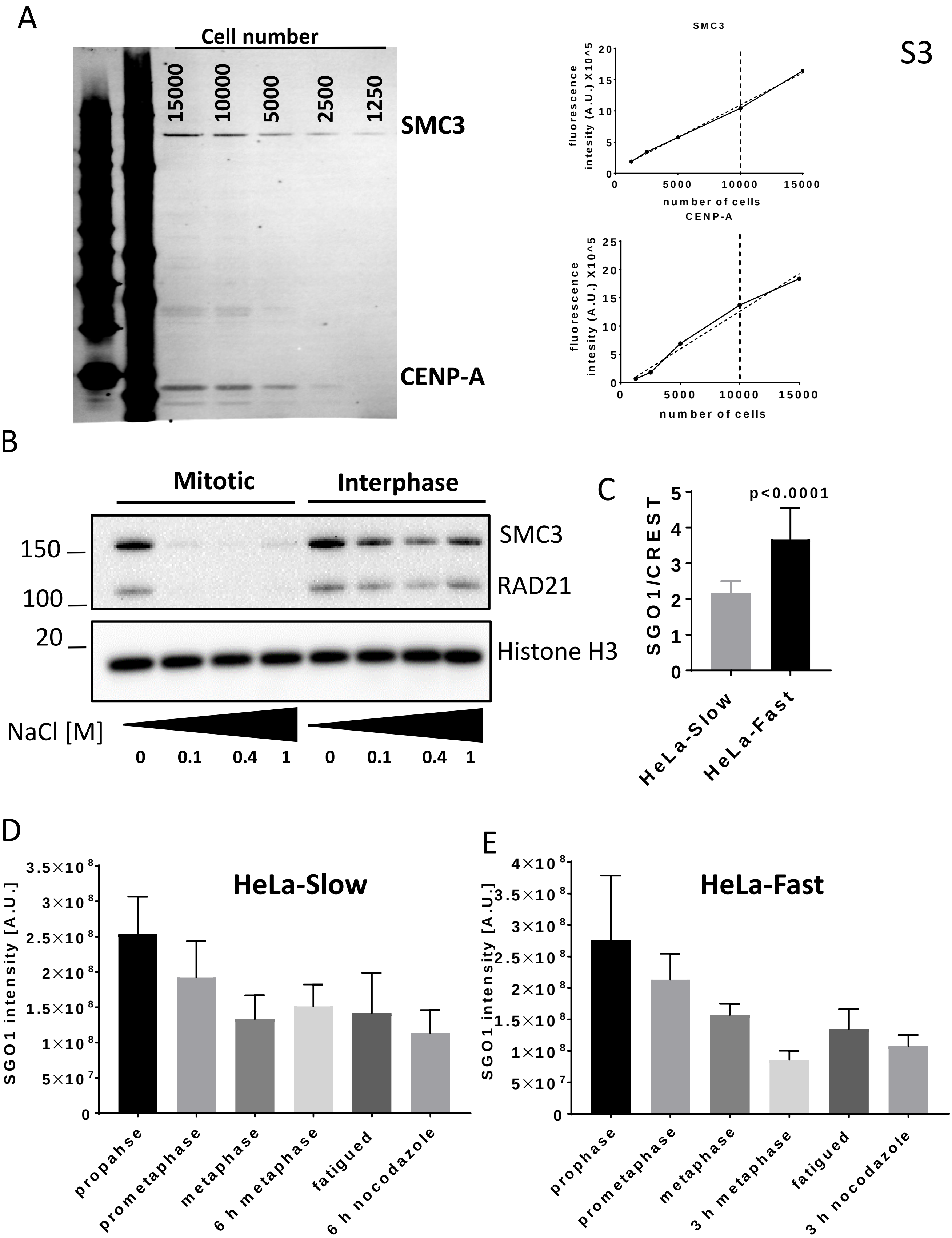

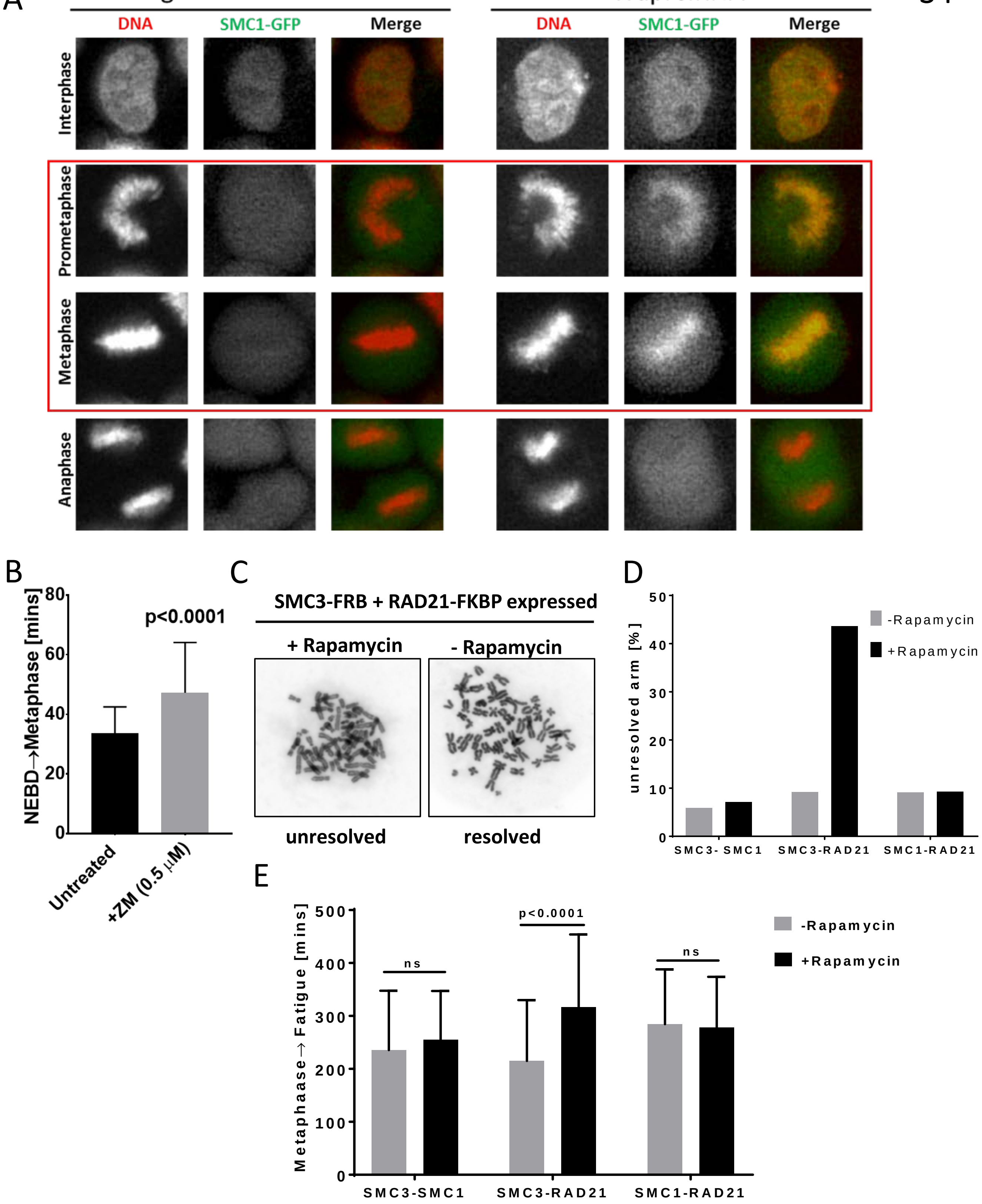

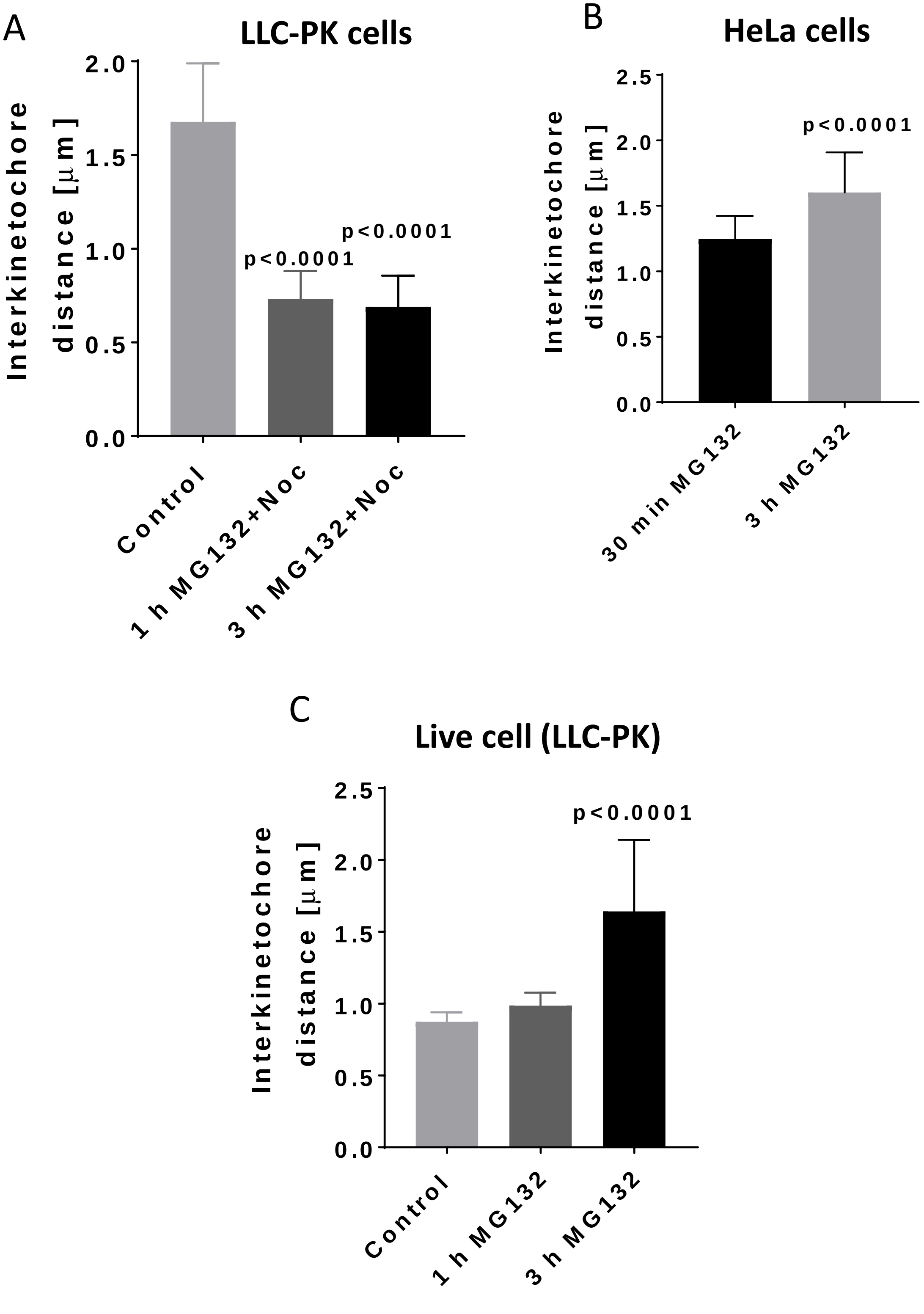

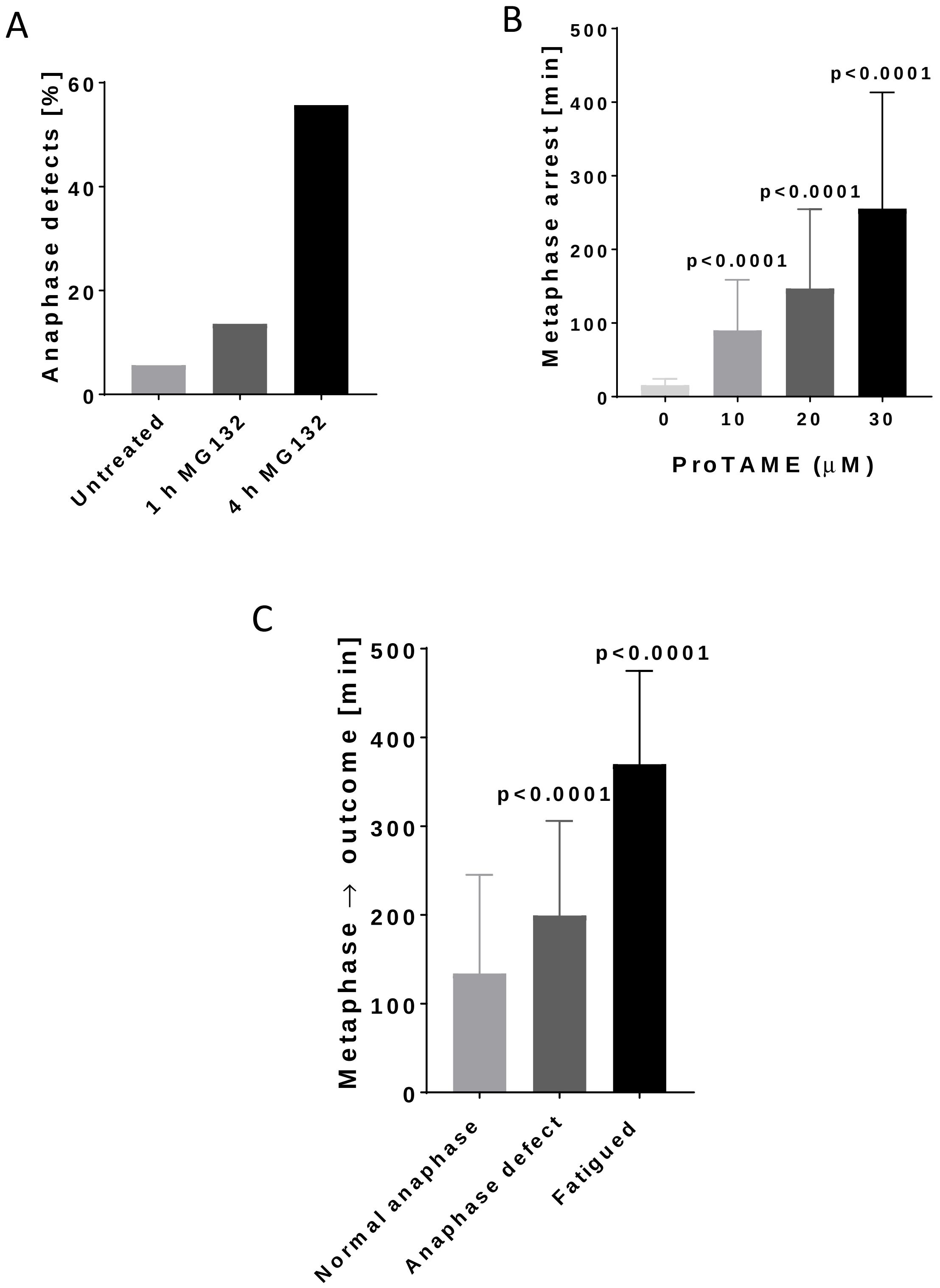
